# Exploring neighborhoods in large metagenome assembly graphs reveals hidden sequence diversity

**DOI:** 10.1101/462788

**Authors:** C. Titus Brown, Dominik Moritz, Michael P. O’Brien, Felix Reidl, Taylor Reiter, Blair D. Sullivan

## Abstract

Genomes computationally inferred from large metagenomic data sets are often incomplete and may be missing functionally important content and strain variation. We introduce an information retrieval system for large metagenomic data sets that exploits the sparsity of DNA assembly graphs to efficiently extract subgraphs surrounding an inferred genome. We apply this system to recover missing content from genome bins and show that substantial genomic sequence variation is present in a real metagenome. Our software implementation is available at https://github.com/spacegraphcats/ spacegraphcats under the 3-Clause BSD License.

---

Metagenomics is the analysis of microbial communities through shotgun DNA sequencing, which randomly samples many subsequences (reads) from the genomic DNA of each microbe present in the community (1).

A common problem in metagenomics is the reconstruction of individual microbial genomes from the mixture. Typically this is done by first running an assembly algorithm that reconstructs longer linear regions based on a graph of the sampled subsequences (2), and then binning assembled contigs together using compositional features and gene content (3, 4). These “metagenome-assembled genomes” are then analyzed for phylogenetic markers and metabolic function. In recent years, nearly 200,000 metagenome-assembled genomes have been published, dramatically expanding our view of microbial life (5–10).

Information present in shotgun metagenomes is often omitted from the binned genomes due to either a failure to assemble (11, 12) or a failure to bin. The underlying technical reasons for these failures include low coverage, high sequencing error, high strain variation, and/or sequences with insufficient compositional or coding signal. Recent work has particularly focused on the problem of strain confusion, in which high strain variation results in considerable fragmentation of assembled content in mock or synthetic communities (11, 12); the extent and impact of strain confusion in real metagenomes is still unknown but potentially significant - metagenome-assembled genomes may be missing 20-40% of true content (13–15).

Associating unbinned metagenomic sequence to inferred bins or known genomes is technically challenging. Some approaches use mapping or k-mer baiting, in which assembled sequences are used to extract reads or contigs from a metagenome or graph (16–20). These methods fail to recover genomic content that does not directly overlap with the query, such as large indels or novel genomic islands. Moreover, most assembly graphs undergo substantial heuristic error pruning and may not contain relevant content (11, 12). Graph queries have shown promise for recovering sequence from regions that do not assemble well but are graph-proximal to the query (21, 22). However, many graph query algorithms are NP-hard and hence computationally intractable in the general case; compounding the computational challenge, metagenome assembly graphs are frequently large, with millions of nodes, and require 10s to 100s of gigabytes of RAM for storage.

In this paper, we develop and implement a scalable graph query framework for extracting unbinned sequence from metagenome assembly graphs with millions of nodes. Crucially, we exploit the structural sparsity of compact De Bruijn assembly graphs in order to compute a succinct index data structure in linear time. Our initial investigations presented here focus on using this index to perform neighborhood queries in large assembly graphs to investigate genome binning and content recovery. This enables us to extract the genome of a novel bacterial species, recover missing sequence variation in amino acid sequences for genome bins, and identify missing content for metagenome-assembled genomes. Our query methods are assembly-free and avoid techniques that may discard strain information such as error correction. These algorithms are available in an open-source Python software package, spacegraphcats (23).

## Results

### Dominating sets enable efficient neighborhood queries in large assembly graphs

We designed and implemented (23) a set of algorithms for efficiently finding content in a metagenome that is close to a query as measured by distance in a compact De Bruijn graph (cDBG) representation of the sequencing data (Figure 1). To accomplish this, we organize the cDBG into *pieces* around a set of *dominators* that are collectively close to all vertices. In this context, the *neighborhood* of a query is the union of all pieces it overlaps; to enable efficient search, we build an invertible index of the pieces.

**Fig. 1.**
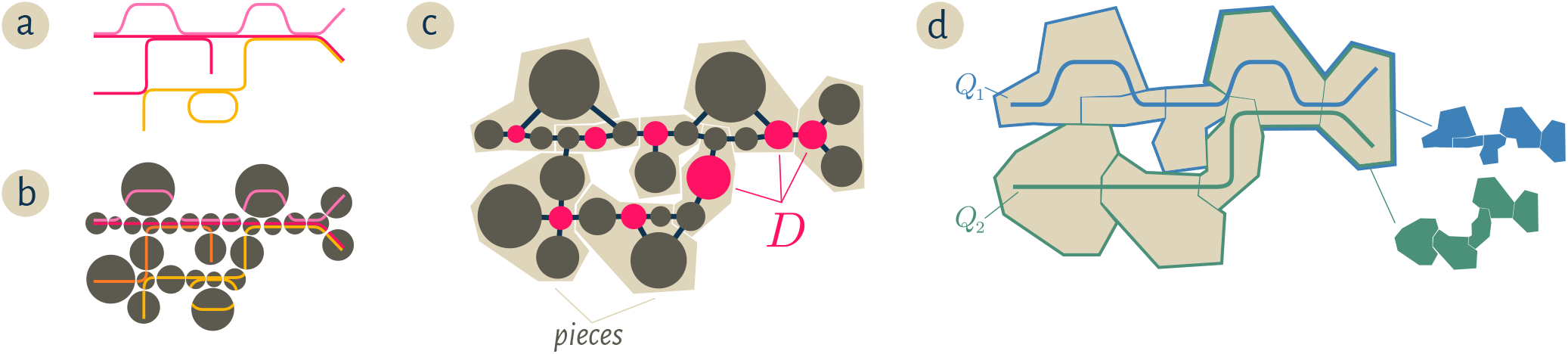
Starting from a collection of genomic sequences (**a**), we form an assembly graph where nodes represent distinct linear subsequences (**b**). In this assembly graph, known as a *compact De Bruijn graph* (4), nodes may represent many k-mers. The original genomic sequences correspond to walks in the graph, and shared nodes between the walks represent shared subsequences. We then (**c**) identify a subset of nodes *D* called a *dominating set* so that every node in the assembly graph is at distance at most one from some member of *D* (marked pink). We further partition the graph into *pieces* by assigning every node to exactly one of the closest members of *D* (beige regions in (**c**) and (**d**)). For a genomic query *Q*, the *neighborhood* of *Q* in this graph is the union of all pieces which share at least one k-mer with the query. The colorful subsets of the pieces in (**d**) correspond to the neighborhoods of the queries *Q*_1_*, Q_2_*.

We compute dominators so that the minimum distance from every vertex in the cDBG to some dominator is at most *r* (an *r-dominating set)* using Algorithm 1, which is based on the linear-time approximation algorithm given by Dvořák and Reidl (24). Although finding a minimum *r*-dominating set is NP-hard (25–27) and an approximation factor below log *n* is probably impossible (26) in general graphs, our approach guarantees constant-factor approximations in linear running time by exploiting the fact that (compact) De Bruijn graphs have *bounded expansion*, a special type of sparsity (28). Algorithm 1 first annotates the graph to determine the distances between all pairs of vertices at distance at most *r* (lines 1-3) and then uses these distances to ensure each vertex is close to a dominator. The core of the efficient distance computation is based on *distance-truncated transitive fraternal (dtf) augmentations* (24) which produce a directed graph 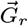 in which each arc *uv* is labeled with *ω*(*uv*), the distance from *u* to *v* in the original cDBG *G*.

Importantly, our implementation enhances the algorithm in (24) by computing only *r*—1 rounds of dtf-augmentations instead of 2*r*—1. Since augmentation is the computationally most expensive part of the pipeline and the running time depends non-linearly on the number of rounds, this vastly improves this algorithm’s scalability. To maintain approximation guarantees on the dominating set size with fewer augmentations, we introduce a threshold parameter `domThreshold`(*r*) which affects the constant factor in our worst-case bound. We selected a threshold (see Supp. Material) that produces *r*-dominating sets of comparable size to those computed by the algorithm in (24). Additionally, we found that processing vertices using a minimum in-degree ordering (line 6) was superior to other common orders (e.g. arbitrary, min/max total degree, max in-degree).

After computing an *r*-dominating set *D* of *G* with Algorithm 1, Algorithm 2 partitions the vertices of *G* into pieces so that each piece *P*[*v*] contains a connected set of vertices for which *v* is a closest member of *D* (Figure 1). Finally, we use minimal perfect hashing (`mphfIndex`) (29) to compute an invertible index^*^ between pieces and their sequence content in the metagenome.

One feature of this approach is that the dominating set and index only need to be computed once for a given metagenome, independent of the number and content of anticipated queries. Queries can then be performed using Algorithm 3 in time that scales linearly with the size of their *neighborhood* – all genomic content assigned to pieces that contain part of the query.

Our implementations of these algorithms in `spacegraphcats` can be run on metagenomic data with millions of cDBG nodes (Table 1); indexing takes under an hour, enabling queries to complete in seconds to minutes (Table 2). See Appendix A for full benchmarking (including cDBG construction). This provides us with a tool to systematically investigate assembly graph neighborhoods.

**Table 1.** Number of cDBG nodes |*V*|, edge density of cDBG |*E*|/|*V*|, size of 1-dominating set |*D*|, average query size (k-mers) 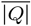, and average number of pieces in query neighborhood 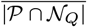. Queries are the 51 genomes and 23 genome bins fully present in `podarV` and `HuSBl`, respectively.

| Dataset | $\vert V\vert $ | $\vert E\vert /\vert V\vert $ | $\vert D\vert $ | $\vert \overline{Q}\vert $ | $\vert \overline{P \cap N_Q}\vert $ |
| --- | --- | --- | --- | --- | --- |
| podarV | 916 041 | 2.2 | 542 350 | 1 475 892 | 4 106 |
| HuSB1 | 13 852 950 | 2.6 | 6 724 505 | 1 112 516 | 106 091 |

**Table 2.** Time and memory usage of spacegraphcats for Algorithms 1-3 on representative metagenome data. The times for Algorithm 3 are averaged over all queries (see Table 1). Statistics reported for Algorithm 2 exclude lines 1-2 of pseudocode. Times are rounded to the nearest tenth of a second; memory is rounded to the nearest Megabyte.

| Dataset | Algorithm | Time (s) | Memory (MB) |
| --- | --- | --- | --- |
| podarV | rdomset | 78.1 | 4 304 |
|  | indexPieces | 359.9 | 14 108 |
|  | search | 14.9 | 3 463 |
| HuSB1 | rdomset | 1 181.1 | 60 238 |
|  | indexPieces | 859.3 | 40 713 |
|  | search | 67.9 | 15 228 |

### Neighborhood queries enable recovery of relevant unknown genomic content

We first measured how well an inexact query can recover a target genome from a metagenome. For a benchmark data set, we used the `podarV` data set (30). This is a “mock” metagenome containing genomes from 65 strains and species of bacteria and archaea, each grown independently and rendered into DNA, then combined and sequenced as a metagenome. This metagenome is a commonly used benchmark for assembly (12, 31–33).

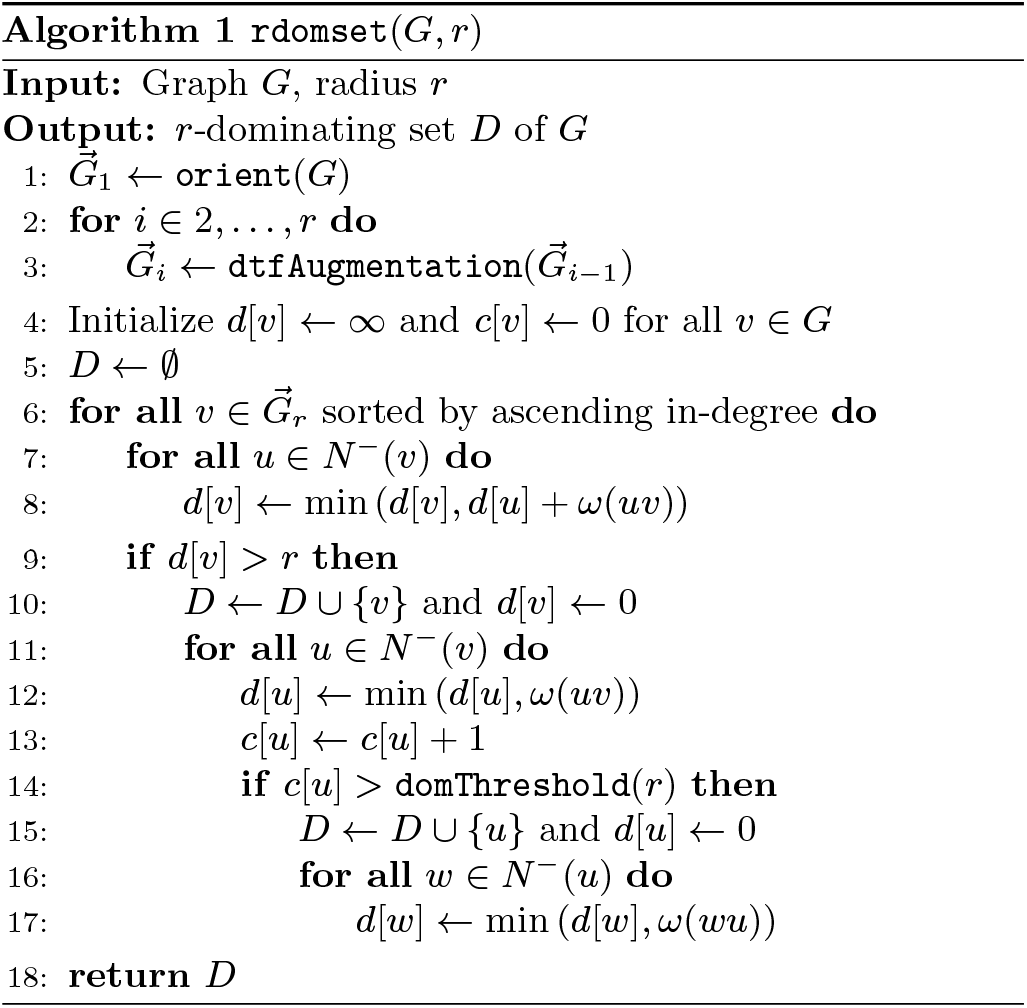

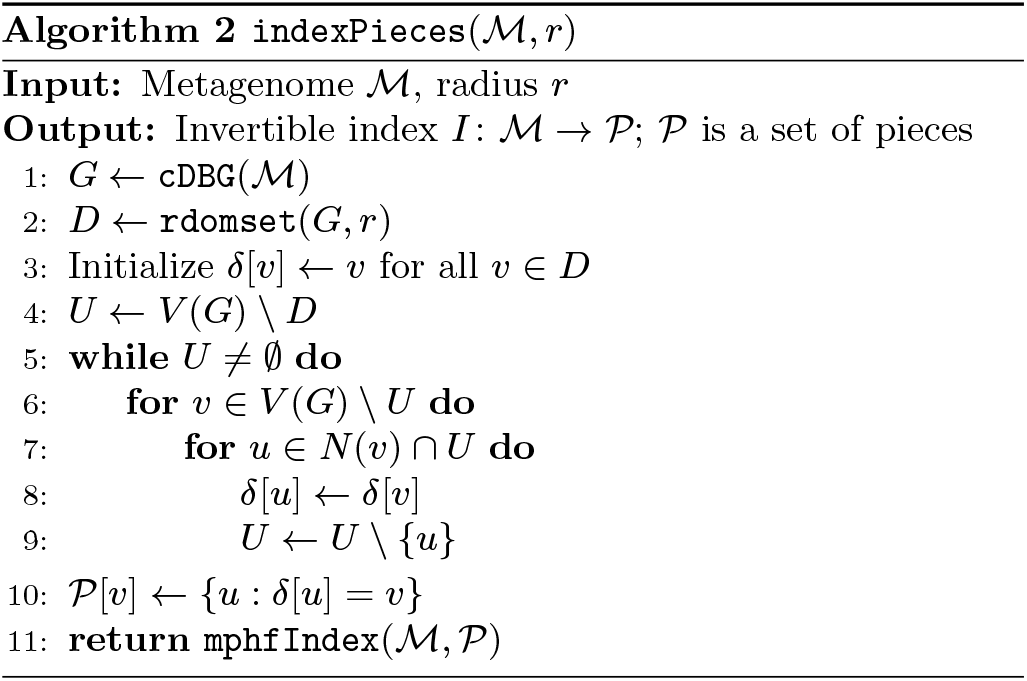

To evaluate the effectiveness of neighborhood query at recovering strain variants, we chose three target genomes from `podarV`– *Porphyromonas gingivalis ATCC 33277, Treponema denticola ATCC 35405*, and *Bacteroides thetaiotaomicron VPI-5482* – that have many taxonomically close relatives in Gen-Bank. We then used these relatives to query the `podarV` mixture and measure the recovery of the target genome. The results, in Figure 2(a), show that graph neighborhood query can recover 35% or more of some target genomes starting from a relative with Jaccard similarity as low as 1%: even a small number of shared k-mers sufficed to define a much larger neighborhood that contains related genomes.

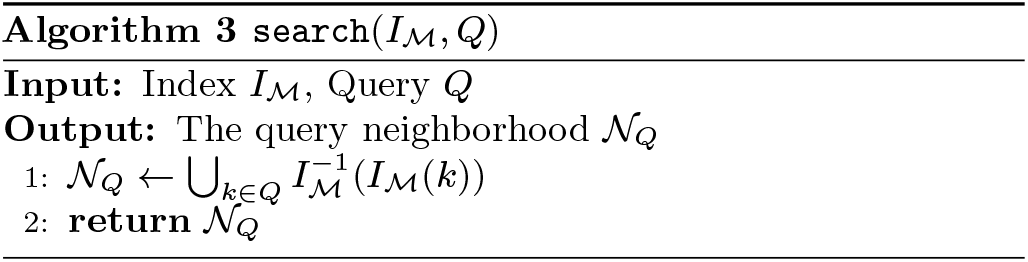

**Fig. 2.**
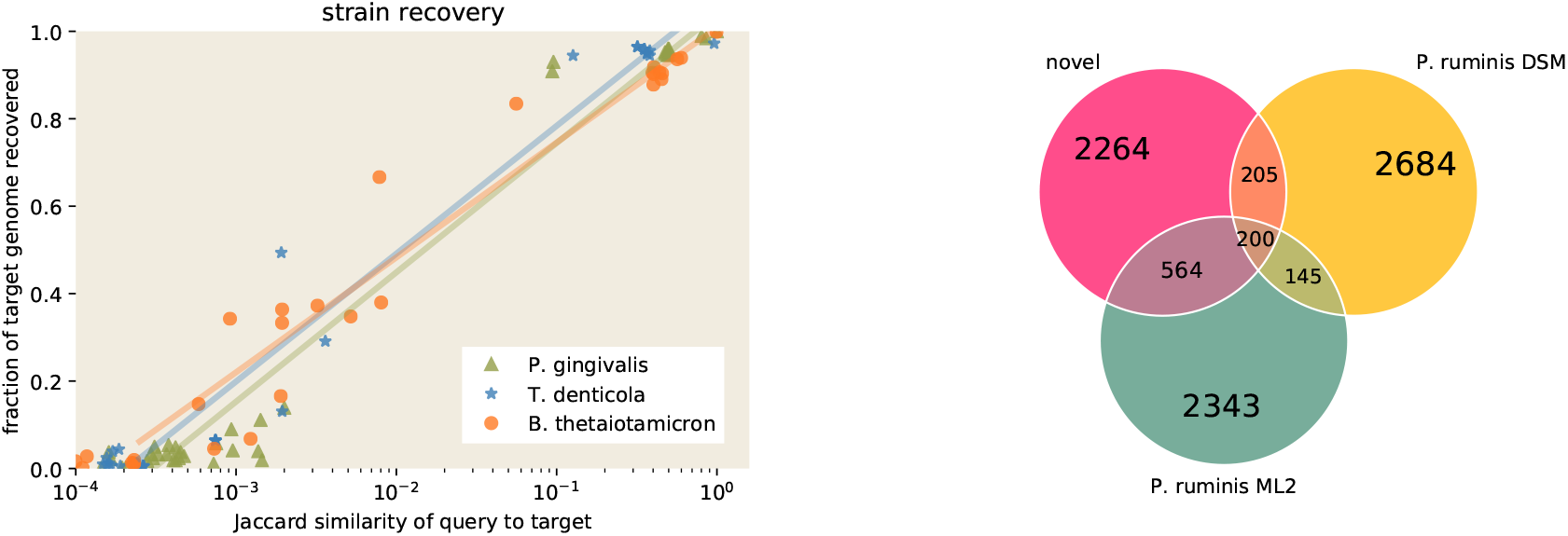
Neighborhood queries enable recovery of relevant genomic content. (a) Left Panel: Recovery of each of three target genomes from `podarV` using queries at a variety of Jaccard distances from the target. Recovery is calculated as containment of target genome in query neighborhood. The solid lines represent logarithmic best-fit curves to the points. (b) Right Panel: Recovery of novel *Proteiniclasticum* content from `podarV`. Nucleotide k-mers from two of the three known *P. ruminis* genomes overlapped approximately a megabase of sequence in the query neighborhood, which also contained approximately 2.3 Mbp of unknown sequence; the third known genome, *P. ruminis CGMCC*, was omitted from the figure as it is 99.7% similar to *P. ruminis DSM*. Numbers are in thousands of k-mers, estimated via sourmash.

We next applied neighborhood query to retrieve an unknown genome from `podarV`. Several papers have noted the presence of unexpected sequence in the assemblies of this data, and Awad et al. identify at least two species that differ from the intended mock metagenome contents (12, 31). One species variant has partial matches to several different *Fusobacterium nucleatum* genomes, while the other incompletely matches to three strains of *Proteiniclasticum ruminis*.

The complete genomes of these two variants are not in public databases and, for the *Proteiniclasticum* variant, cannot be entirely recovered with existing approaches: when we assemble the reads that share k-mers with the available genomes, a marker-based analysis with CheckM estimates that 98.8% of the *Fusobacterium* variant is recovered, while only 72.96% of the *Proteiniclasticum* variant is recovered. We therefore tried using neighborhood queries to expand our knowledge of the *Proteiniclasticum* variant.

We performed a neighborhood query into `podarV` with all three known *Proteiniclasticum* genomes from GenBank. We then extracted the reads overlapping this neighborhood and assembled them with MEGAHIT. The retrieved genome neighborhood for *Proteiniclasticum* contains 2264K novel k-mers (Figure 2(b)). The reads from the query neighborhood assembled into a 3.1 Mbp genome bin. The assembly is estimated by CheckM to be 100% complete, with 10.3% contamination. The mean amino acid identity between *P. ruminis ML2* and the neighborhood assembly is above 95%, suggesting that this is indeed the genome of the *Proteiniclasticum* variant, and that neighborhood query retrieves a full draft genome sequence (see Supp. Material A).

### Query neighborhoods in a real metagenome do not always assemble well

Real metagenomes may differ substantially from mock metagenomes in size, complexity, and content. In particular, real metagenomes may contain a complex mixture of species and strain variants (34) and the performance of assembly and binning algorithms on these data sets is challenging to evaluate in the absence of ground truth. One recent comparison of single-cell genomes and metagenome-assembled genomes in an ocean environment found that up to 40% of single-cell genome content may be missing in metagenome-assembled genomes (15).

We first ask whether genome query neighborhood sizes in a real metagenome differ from mock metagenomes. We examined genomes inferred from the SB1 sample from the Hu et al. (2016) study, in which 6 metagenomic samples taken from various types of oil reservoirs were sequenced, assembled, binned, and computationally analyzed for biochemical function (35). Examining the 23 binned genomes in GenBank originating from the SB1 sample, we compared the `HuSB1` neighborhood size distribution with the `podarV` data set (Figure 3(a)). We saw that more genome bins in `HuSB1` have 1.5x or larger query neighborhoods than do the genomes in `podarV`. This suggests the presence of considerably more local neighborhood content in the real metagenome than in the mock metagenome.

**Fig. 3.**
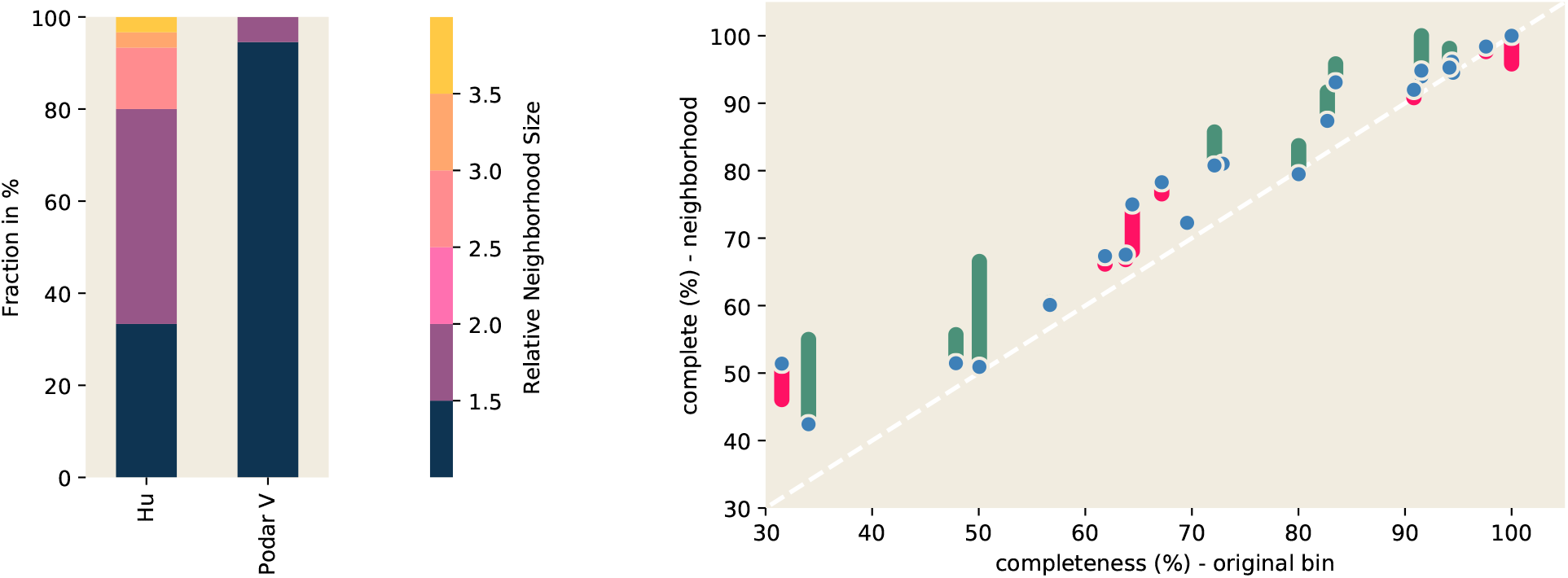
Query neighborhoods in `HuSBl` metagenome are large and contain additional marker genes. (a) Left Panel: Neighborhood sizes are larger in `HuSBl` than in `podarV`. Here we queried `podarV` and `HuSBl` using each of 51 and 23 genomes fully present in the respective datasets and measured the relative size of its neighborhood—a size of 1 indicates that no additional sequence is present in the neighborhood, while a size of 2 indicates that the retrieved neighborhood is twice the size of the query genome. (b) Right Panel: Query neighborhoods are estimated to be more complete than the original genome bins. We queried `HuSBl` using each of 23 genomes binned from SB1, and assembled the resulting neighborhoods using MEGAHIT and Plass. The blue points represent completeness estimates of MEGAHIT-assembled neighborhoods, while green and pink bars represent the additional or missing content in the Plass assemblies respectively.

We next investigated metagenomic content in the query neighborhoods. As with the unknown variants in `podarV`, we used CheckM to estimate genome bin completeness. The estimated bin completeness for many of the query genomes is low (Appendix A). To see if the query neighborhoods contain missing marker genes, we assembled reads from the query neighborhoods using MEGAHIT, and found this improved the completion metrics (Figure 3(b)).

Investigating further, we found that the query neighborhood assemblies contain only between 4% and 56% of the neighborhood k-mer content (Appendix A), suggesting that MEGAHIT is not including many of the reads in the assembly of the query neighborhoods. This could result from high error rates and/or high strain variation in the underlying reads (11, 12).

To attempt the recovery of more gene content from the assemblies, we turned to the Plass amino acid assembler (36). Plass implements an overlap-based amino acid assembly approach that is considerably more sensitive than nucleotide assemblers and could be more robust to errors and strain variation (37).

When we applied Plass to the reads from the query neighborhoods, we saw a further increase in neighborhood completeness (Figure 3(b)). This suggests that the genome bin query neighborhoods contain real genes that are accessible to the Plass amino acid assembler. We note that these are unlikely to be false positives, since CheckM uses a highly specific Hidden Markov Model (HMM)-based approach to detecting marker genes (38).

### Some query neighborhoods contain substantial strain variation

If strain variation is contributing to poor nucleotide assembly of marker genes in the query neigborhoods, then Plass should assemble these variants into similar amino acid sequences. Strain variation for unknown genes can be difficult to study due to lack of ground truth, but highly conserved proteins should be readily identifiable.

The *gyrA* gene encodes an essential DNA topoisomerase that participates in DNA supercoiling and was used by (35) as a phylogenetic marker. In the GenBank bins, we found that 15 of the 23 bins contain at least one gyrA sequence (with 18 *gyrA* genes total). We therefore used *gyrA* for an initial analysis of the Plass-assembled neighborhood content for all 23 bins. To avoid confounding effects of random sequencing error in the analysis and increase specificity at the cost of sensitivity, we focused only on high-abundance data: we truncated all reads in the query neighborhoods at any k-mer that appears fewer than five times, and ran Plass on these abundance-trimmed reads from each neighborhood. We then searched the gene assemblies with a gyrA-derived HMM, aligned all high-scoring matches, and calculated a pairwise similarity matrix from the resulting alignment.

When we examine all of the high-scoring gyrA protein matches in the hard-trimmed data, we see considerable sequence variation in some query neighborhoods (Figure 4(a)). Much of this variation is present in fragmented Plass assemblies; when the underlying nucleotide sequences are retrieved and used to construct a compact De Bruijn graph, the variation is visible as spurs off of a few longer paths (insets in Figure 4(a)). When we count the number of well-supported amino acid variants in isolated positions (i.e. ignoring linkage between variants) we see that ten of the 23 neighborhoods have an increased number of gyrA genes, with four neighborhoods gaining a gyrA where none exists in the bin (Appendix A; see lowest inset in Figure 4(a) for one example). Only one neighborhood, *M. bacterium*, loses its gyrA genes due to the stringent k-mer abundance trimming. Collectively, the use of the Plass assembler on genome neighborhoods substantially increases the number of gyrA sequences associated with bins.

**Fig. 4.**
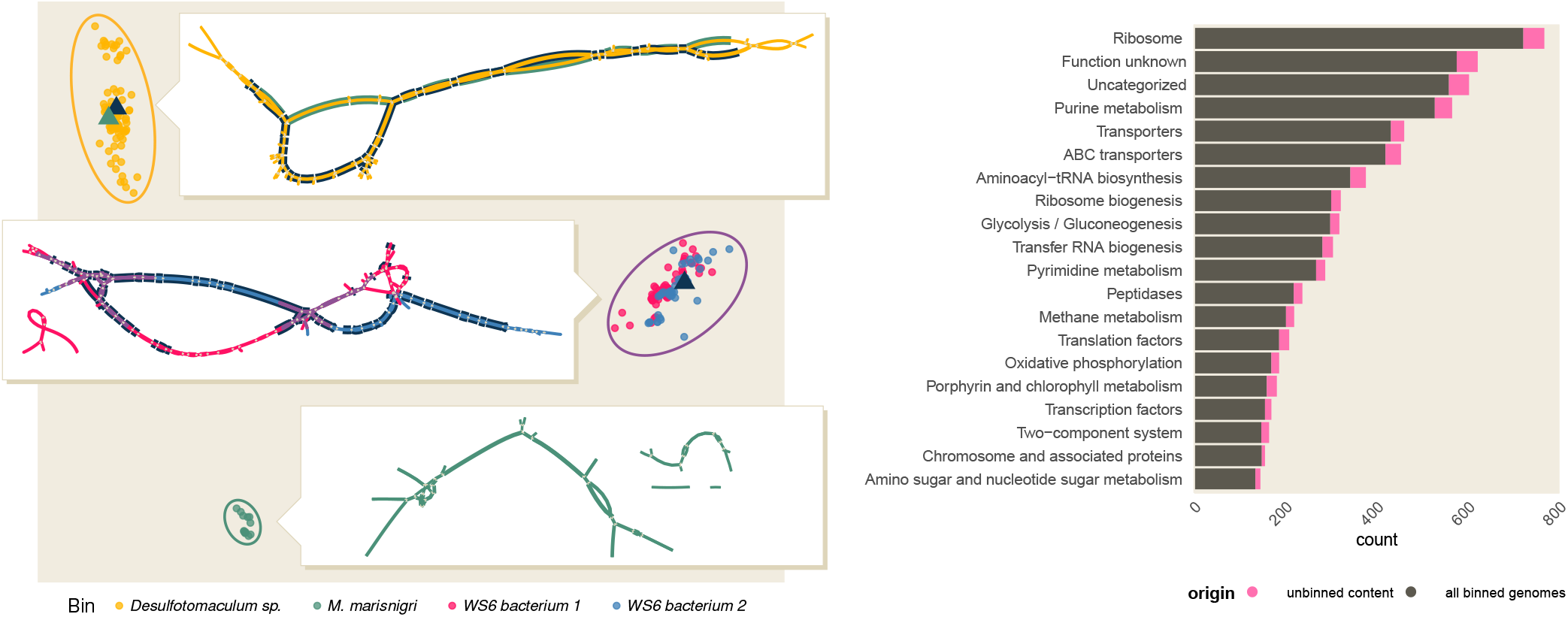
Query neighborhoods in `HuSBl` contain sequence variants and new genes. (a) Left Panel: gyrA has substantial minor sequence variation in several query neighborhoods. In this multidimensional scaling plot, each point represents a distinct gyrA sequence from the Plass assemblies of four representative query neighborhoods, colored by query binned genome. The triangles represent gyrA sequences originating from the query binned genome, if any are present. The inlays are visualizations of assembly graphs of reads that contain *gyrA* sequence in each neighborhood. Unitigs are colored by their cluster of origin; matches to *gyrA* sequences from the bin are highlighted using color from relevant triangle. (b) Right Panel: Genome neighborhoods re-associate annotated functionality to binned genomes. For each of 23 genome bins originating from `HuSBl`, we found the unbinned content by removing all orthologs found in the binned genomes in (39) and by counting distinct ortholog annotations once. Functional content is distributed throughout pathways present in the binned genomes, and increases functionality associated with binned genomes by approximately 13%.

We see this same pattern for many genes, including *alaS*, *gyrB, rpb2* domain 6, *recA, rplB*, and *rpsC* (Appendix A). This shows that multiple variants of those proteins are present within at least some of the neighborhoods and implies the presence of underlying nucleotide strain variation. This strain variation may be one reason that nucleotide assembly performs poorly: on average, only 19.6% of Plass-assembled proteins are found within the nucleotide assemblies.

### Query neighborhoods assembled with Plass contain additional functional content

In addition to capturing variants close to sequences in the bins, we identify many novel genes in the query neighborhoods. We used KEGG to annotate the Plass-assembled amino acid sequences, and then removed any annotations already present for genes in the genome bin. We also ignored homolog abundance such that each homolog is counted only once per neighborhood.

Novel functional content is distributed throughout pathways present in the genome bins, and increases functionality associated with binned genomes by approximately 13% (Figure 4(b)). This includes orthologs in biologically relevant pathways such as methane metabolism, which are important for biogeochemical cycling in oil reservoirs (35).

Genes in these neighborhoods contain important metabolic functionality expanding the pathways already identified in (35). We find 40 unique orthologs involved in nitrogen fixation across eight neighborhoods, four of which had no ortholog in the bin. Importantly, we find the ratio of observed orthologs approximately matches that noted in (35), where two thirds of nitrogen fixation functionality is attributable to archaea (29 of 40 orthologs). This is in contrast to most ecological systems where bacteria are the dominant nitrogen fixers (35).

## Discussion

### Efficient graph algorithms provide novel tools for investigating graph neighborhoods

Recent work has shown that incorporating the structure of the assembly graph into the analysis of metagenome data can provide a more complete picture of gene content (21, 22). While this has provided evidence that it is useful to analyze sequences that have small graph distance from a query (are in a “neighborhood”), this approach has not been widely adopted. Naïvely, local expansion around many queries in the assembly graph does not scale to these types of analyses due to the overhead associated with searching in a massive graph. The neighborhood index structure described in this work overcomes this computational obstacle and enables rapid exploration of sequence data that is local to a query.

Because a partition into pieces provides an implicit data reduction (the cDBG edge relationships are subsumed by piece membership), the query-independent nature of the index allows many queries to be processed quickly without loading the entire graph into memory. Our approach consequently provides a data exploration framework not otherwise available.

Exploiting the structural sparsity of cDBGs is a crucial component of our algorithms. First, it is necessary to use graph structure to obtain a guarantee that Algorithm 2 finds a small number of pieces since the size of a minimum *r*-dominating set cannot be approximated better than a factor of log *n* in general graphs^†^ unless NP ⊆ DTIME(*n*^*O*(log log *n*)^) (26). Without such a guarantee, we cannot be sure that we are achieving significant data reduction by grouping cDBG vertices into pieces. Being able to do this in linear time also ensures that indexing and querying can scale to very large data sets. Furthermore, because we utilize a broad structural characterization (bounded expansion) of cDBGs rather than a highly specialized aspect, our methods enable neighborhood-based information retrieval in any domain whose graphs exhibit bounded expansion structure; examples include some infrastructure, social, and communication networks (24, 40, 41).

### Neighborhood queries extend genome bins

In both the `podarV` and `HuSBl` metagenomes, neighborhood queries were able to identify additional content likely belonging to query genomes. In the `podarV` mock metagenome, we retrieved a potentially complete genome for an unknown strain based on partial matches to known genomes. In the `HuSBl` metagenome, we increased the estimated completeness of most genome bins – in some cases substantially, e.g. in the case of *P_bacterium 34_609* we added an estimated 20.9% to the genome bin. In both cases we rely solely on the structure of the assembly graph to expand the genome bins. We do not make use of sequence composition, contig abundance, or phylogenetic marker genes in our search. Thus graph proximity provides an orthogonal set of information for genome-resolved metagenomics that could be used to improve current binning techniques.

### Query neighborhoods from real metagenomes contain substantial strain variation that may block assembly

Previous work suggests that metagenome assembly and binning approaches are fragile to strain variation (11, 12). This may prevent the characterization of some genomes from metagenomes. The extent of this problem is unknown, although the majority of approaches to genome-resolved metagenomics rely on assembly and thus could be affected.

In this work, we find that some of the sequence missing from genome bins can be retrieved using neighborhood queries. For `HuSBl`, some genome bins are missing as many as 68.5% of marker genes from the original bins, with more than half of the 22 bins missing 20% or more; this accords well with evidence from a recent comparison of single-cell genomes and metagenome-assembled genomes (15), in which it was found that metagenome-assembled genomes were often missing 20% to 40% of single-cell genomic sequence. Neighborhood query followed by amino acid assembly recovers additional content for all but two of the genome bins; this is likely an underestimate, since Plass may also be failing to assemble some content.

When we bioinformatically analyze the function of the expanded genome content from neighborhood queries, our results are consistent with the previous metabolic analyses by (35), and extend the set of available genes by 13%. This suggests that current approaches to genome binning are specific, and that the main question is sensitivity, which agrees with a more direct measurement of lost content (15).

### Neighborhood queries enable a genome-targeted workflow to recover strain variation

The spacegraphcats analysis workflow described above starts with genome bins. The genome bins are used as a query into the metagenome assembly graph, following which we extract reads from the query neighborhood. We assemble these reads with the Plass amino acid assembler, and then analyze the assembly for gene content. We show that the Plass assembly contains strain-level heterogeneity at the amino acid level, and that this heterogeneity is supported by underlying nucleotide diversity. Even with stringent error trimming on the underlying reads, we identify at least thirteen novel gyrA sequences in ten genome neighborhoods.

Of note, this workflow explicitly associates the Plass assembled proteins with specific genome bins, as opposed to a whole-metagenome Plass assembly which recovers protein sequence from the entire metagenome but does not link those proteins to specific genomes. The binning-based workflow connects the increased sensitivity of Plass assembly to the full suite of tools available for genome-resolved metagenome analysis, including phylogenomic and metabolic analysis. However, spacegraphcats does not separate regions of the graph shared in multiple query neighborhoods; existing strain recovery approaches such as DESMAN or MSPminer could be used for this purpose (16, 19).

One future step could be to characterize unbinned genomic content from metagenomes by looking at Plass-assembled marker genes in the metagenome that do not belong to any bin’s query neighborhood. This would provide an estimate of the extent of metagenome content remaining unbinned.

## Conclusions

The neighborhood query approach described in this work provides an alternative window into metagenome content associated with binned genomes. We extend previous work showing that assembly-based methods are fragile to strain variation, and provide an alternative workflow that substantially broadens our ability to characterize metagenome content. This first investigation focuses on only two data sets, one mock and one real, but the neighborhood indexing approach is broadly applicable to all shotgun metagenomes.

In this initial investigation of neighborhood indexing, we have focused on using neighborhood queries with a genome bin. We recognize that this approach is of limited use in regions where no genome bin is available; spacegraphcats is flexible and performant enough to support alternative approaches such as querying with k-mers belonging to genes of interest.

Potential applications of spacegraphcats in metagenomics include developing metrics for genome binning quality, analyzing pangenome neighborhood structure, exploring *r*-dominating sets for *r* > 1, extending analyses to colored De Bruijn graphs, and investigating *de novo* extraction of genomes based on neighborhood content. We could also apply spacegraphcats to analyze the neighborhood structure of assembly graphs overlayed with physical contact information (from e.g. HiC), which could yield new applications in both metagenomics and genomics (42, 43).

More generally, the graph indexing approach developed here may be applicable well beyond metagenomes and sequence analysis. The exploitation of bounded expansion to efficiently compute *r*-dominating sets on large graphs makes this technique applicable to a broad array of problems.

## Materials and Methods

### Data

We use two data sets: SRR606249 from `podarV` (44) and SRR1976948 (sample SB1) from `hu` (39). Each data set was first preprocessed to remove low-abundance k-mers as in (45), using `trim-low-abund.py` from khmer v2.1.2 (46) with the parameters `-C 3 -Z 18 -M 20e9 -V -k 31`. We build compact De Bruijn graphs using BCALM v2.2.0 (47). Stringent read trimming at low-abundance k-mers was done with `trim-low-abund.py` from khmer, with the parameters `-C 5 -M 20e9 -k 31`.

### Software

The source code for the index construction and search is available at https://github.com/spacegraphcats/spacegraphcats (23). It is implemented in Python 3 under the 3-Clause BSD License. Version 1.1, used in this paper, is archived at DOI: 10.5281/zenodo.2505206.

Snakemake (48) workflows to reproduce all of the analysis are available at https://github.com/spacegraphcats/2018-paper-spacegraphcats/, and Jupyter Notebooks to recreate Figures 1, 2, 3, and 4(b) are in that same repository (49). The notebooks rely on the numpy, matplotlib, pandas, scipy, and Vega-Lite libraries (50–54). The workflow repository is archived at DOI: 10.5281/zenodo.2592780.

### Benchmarking

We measured time and memory usage for Algorithms 1-3 by executing the following targets in the spacegraphcats `conf/Snakefile: catlas.csv` for `rdomset, contigs.fa.gz.mphf` for `indexPieces`, and `search` for `search`. We report wall time and maximum resident set size, running under Ubuntu 18.04 on an NSF Jetstream virtual machine with 10 cores and 30 GB of RAM (55, 56). To measure maximum resident set size, we used the memusg script (Jaeho Shin, https://gist.github.com/netj/526585).

### Graph denoising

For each data set, we built a compact De Bruijn graph (cDBG) for k=31 by computing the set of unitigs with BCALM (57) and removing all vertices of degree one with a mean *k*-mer abundance of 1.1 or less. After the removal of these vertices, we then contracted any newly revealed degree-two paths.

### Neighborhood indexing and search

We used spacegraphcats to build an *r*-dominating set for each denoised cDBG and index it. We then performed neighborhood queries with spacegraphcats, which produces a set of cDBG nodes and reads that contributed to them. The full list of query genomes for the *Proteiniclasticum* query is available in Supp Material A, and the NCBI accessions for the *P. gingivalis, T. denticola*, and *B. thetaiotamicron* queries are in the directory `pipeline-base` of the paper repository, files `strain-gingivalis.txt, strain-denticola.txt`, and `strain-bacteroides.txt`, respectively.

### Search results analysis

Query neighborhood size, Jaccard containment, and Jaccard similarity were estimated using modulo hash signatures with a k-mer size of 31 and a scaled factor of 1000, as implemented in sourmash v2.0a9 (58).

### Assembly and genome bin analysis

We assembled reads using MEGAHIT v1.1.3 (31) and Plass v2-c7e35 (36), treating the reads as single-ended. Bin completeness was estimated with CheckM 1.0.11, with the `–reduced_tree` argument (38). Amino acid identity between bins and genomes was calculated using CompareM commit 7cd51276 (https://github.com/dparks1134/CompareM).

### Gene targeted analysis

Analysis of specific genes was done with HMMER v3.2.1 (59). Plass amino acid assemblies were queried with `HMMER hmmscan` using the PFAM domains listed in Supp Material 9, using a threshold score of 100 (60). Matching sequences were then extracted from the assemblies for further analysis. To overcome problems associated with comparing non-overlapping sequence fragments, only sequences that overlapped 125 of the most-overlapped 200 residues of the PFAM domain were retained (all sequences shared a minimum overlap of 50 residues with all other sequences). These sequences were aligned with MAFFT v7.407 with the `–auto` argument (61). Pairwise similarities were calculated using HMMER where the final value represented the number of identical amino acids in the alignment divided by the number of overlapping residues between the sequences. Pairwise distances were visualized using a multidimensional scaling calculated in R using the `cmdscale` function. To visualize the assembly graph structure underlying these amino acid assemblies, we used paladin v1.3.1 to map abundance-trimmed reads back to the Plass amino acid assembly, with `-f 125` to set the minimum ORF length accepted (62). We extracted the reads that mapped to the gene of interest, created an assembly graph using BCALM v2.2.0 (57), and visualized the graph using Bandage v0.8.1 (63). We colored nucleotide sequences originating from the bins using the BLAST feature in Bandage.

### KEGG Analysis

We annotated the Plass assemblies using Kyoto Encyclopedia of Genes GhostKOALA v2.0 (64). To assign KEGG ortholog function, we used methods implemented at https://github.com/edgraham/GhostKoalaParser release 1.1.

## ACKNOWLEDGMENTS

This work is funded in part by the Gordon and Betty Moore Foundation’s Data-Driven Discovery Initiative through Grants GBMF4551 to C. Titus Brown, GBMF4553 to Jeffrey Heer, and GBMF4560 to Blair D. Sullivan. This work arose from the Barnraising for Data-Intensive Discovery at Mt. Desert Island Biological Lab in May 2016. We thank Erich Schwarz, Martin Steinegger, Johannes Söding, Mark Blaxter, and members of the Data Intensive Biology lab at UC Davis for discussion and feedback.

## Appendix

### Notation

For a graph *G* we write *V*(*G*) and *E*(*G*) for its vertex and edge set, respectively. For a vertex *v* ∈ *V*(*G*), we write *N*(*v*) to denote its neighborhood, i.e. the set containing all vertices that are adjacent to it in *G*. We further write *G* – *v* for the graph obtained from *G* by removing *v* and all edges incident to it.

For a directed graph (digraph) 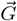, we use *N*^−^ (*v*) to denote the *in-neighborhood* of 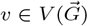, that is, the set of all vertices *w* for which an arc *wv* exists in 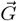. The size of that in-neighbourhood is called the *in-degree* of *v*. We write 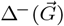 for the maximum in-degree over all vertices of 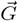. For ease of notation, we usually write “*v* ∈ *G*” instead of “*v* ∈ *V* (*G*)” and “*uv* ∈ *G*” instead of “{*u,v*} ∈ *E*(*G*)” (same for digraphs).

A vertex set *S* ⊆ *V*(*G*) is *r-scattered* if all vertices in it have pairwise distance at least *r* in *G*. Note that if a set *S* is (2*r* + 1)-scattered in *G*, then no single vertex can *r*-dominate more than one vertex of *S*. It follows that any *r*-dominating set of *G* must have size at least |*S*|; we can therefore use (2*r* + 1)-scattered sets to prove lower bounds on the *r*-domination number of a graph.

### Dtf-augmentation

For the sake of completeness, we briefly describe how dtf-augmentations 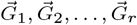 are computed for an input graph *G*.

First, the graph 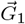 is a *minimum in-degree orientation* of *G* obtained as follows: we find a vertex *u* of minimum degree in *G*, orient the incident edges towards *u* and the repeat the procedure in *G* – *u* until no vertices remain.

Given 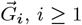 we then compute 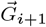 by applying an *augmentation step*:

1. 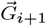 contains all arcs of 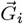,
2. if there is a pair of arcs 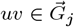 and 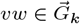 with *j* + *k* = *i* + 1, then we add *uw* to 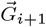,
3. if there is a pair of arcs 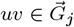 and 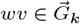 with *j* + *k* = *i* + 1, then either the arc *uw* or the arc *wu* must be in 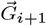.

The ambiguity in the last step is resolved as follows: we collect all edges *uw* for which the last case applies in a graph *H* and then compute a minimum in-degree orientation 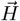 of *H*. We then add the arcs in 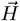 to 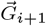.

For ease of notation, we define *ω*(*uv*) to be the smallest index *i* ≤ *r* such that 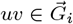 (and ∞ if no such index exists). We make use of two crucial properties of dtf augmentations in the following. First, vertices that have distance at most *r* in *G* have distance at most two in 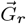:

#### Lemma 1

(c.f. Lemma 26 in (24)). *For every pair u,v ∈ G with dist(u, v) = d and for every integer r ≥ d one of the following holds:*

1. 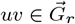 *and ω*(*uv*) = *d*,
2. 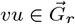 *and ω*(*vu*) = *d*,
3. *or there exists x such that xu*, 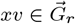 *and ω*(*xu*) + *ω*(*xv*) = *d*.

Second, if we compute dtf-augmentations of graphs from a bounded expansion class (to which graph of bounded maximum degree like cDBGs belong), then the maximum in-degree of its *r*th augmentation can be bounded by a function independent of the *size* of the graph:

#### Theorem 1

(c.f. Theorem 16 in (24)). *For every graph class* 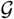 *with bounded expansion there exists a function f such that for every member* 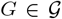 *the rth dtf-augmentation* 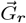 *computed as above satisfies* 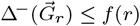.

#### Approximation guarantee

Let us introduce some notation for the analysis of Algorithm 1. We first partition the vertices of *D* according to whether they were added in line 10 (denoted by *D*_1_) or in line 15 (denoted by *D*_2_). Let *v*_1_,…,*v_n_* be the vertex-order in which they are iterated over in the loop starting at line 6. We will use the notation 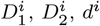, and *c^i^* to represent the states of the respective sets and data structures during the *i*th iteration of said loop. Let *τ*:= domThreshold (*r*) be the chosen threshold (we discuss a good value for *τ* on cDBGs below).

##### Lemma 2.

*After the for-loop at line 7 has finished*,

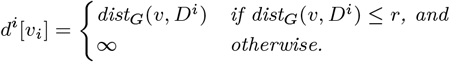

*Proof*. The statement trivially holds while *D^z^* = 0, so assume otherwise. Let *u_h_* ∈ *D^i^* be the vertex closest to *v_i_* and let *h* < *i* be the iteration in which *u_h_* was added to *D* (either in line 10 or line 15 of that iteration).

If *d*:= dist_*G*_(*v_i_,u_h_*) > *r*, then *d^i^*[*v_i_*] has not been changed yet and is still set to ∞. Otherwise, consider the three possible scenarios promised by the distance-property of dtf-augmentations:

##### Case 1

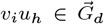. Then *ω*(*v_i_u_h_*) = *d* and in iteration *h* the value of *d^h^*[*v_i_*] is set to the correct value *d* at line 8. By assumption this distance remains minimal until iteration *i* and hence *d^i^*[*v_i_*] = *d^h^*[*v_i_*] = *d*.

##### Case 2

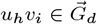. Then *ω*(*v_i_u_h_*) = *d* and in iteration *i* the value of *d^i^*[*v_i_*] is set to the correct value *d* at line 8.

##### Case 3

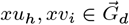 with *ω*(*xu_h_*) + *ω*(*xv_i_*) = *d*.

During iteration *h* the value of *d^h^*[*x*] is set to *ω*(*xu_h_*) at line 8 and subsequently retrieved in iteration *i* when *d^i^*[*v_i_*] is set to

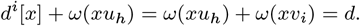

We conclude that after the execution of the loop at line 8. *d^i^*[*v_i_*] is set to ∞ if *v_i_* is not dominated by *D_i_* and is otherwise set to dist_*G*_(*v_i_, D^z^*), as claimed.

As an immediate consequence, we see the conditional statement at the end of the loop at line 8 accurately determines whether *v_i_* is dominated by *D_i_* or not. Accordingly, line 15 of the loop is only executed if *v_i_* is *not* dominated by *D^i^*. Another consequence is that all vertices in *D*_1_ have large distance to each other:

##### Corollary 1.

*The set D*_1_ *is* (*r* + 1)-*scattered in G*.

We need one more important property of the algorithm in order to derive the approximation factor.

##### Lemma 3.

*For every w ∈ G it holds that* 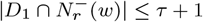.

*Proof*. Assume towards a contradiction that *τ* + 2 such vertices *v*_*i*_1__,…, *v_i_τ_ + 2_, i*_1_ < *i*_2_ < … < *i*_*τ*+2_ exist in 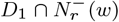. Since every such vertex *v_i_, i* ∈ {*i*_1_,… *i*_*τ*+2_}, was added to *D* in part (2), part (3) of the algorithm was executed during iteration *i* as well. Thus *c*[*w*] was increased in each iteration *i* and during iteration *i*_*τ*+1_ we have that *c*[*w*] ≥ *τ* + 1 after the increment of *c*[*w*]. Therefore part (4) must have been executed for *w*, including *w* into *D*. Hence *w* ∈ *D^s^* for *s* > *i*_*τ*+1_ and in particular *w ∈ D*^*i*_*τ*+2_^. But then *v*_*i*_*τ*+2__ was dominated by *w* at the beginning of iteration *i*_*τ*+2_ since we assumed that *ω*(*rv*_*i*_*τ*+2__) ≤ *r*, thus *v*_*i*_*τ*+2__ would not have been included in *D* at step (2). This contradicts our assumption of *v*_*i*_*τ*+2__ ∈ *D*_1_ so the claim must hold.

##### Lemma 4.

*There exists a subset A ⊆ D*_1_ *such that A is* (2*r* +1)- *scattered in G and*

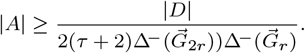

*Proof*. We construct an auxiliary graph *H* with vertices *D*_1_ by adding arcs *v_i_v_j_* for *v_i_, v_j_* ∈ *D*_1_ with *i* < *j* whenever dist_*G*_(*v_i_, v_j_*) ≤ 2*r*. Let 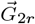 be a 2*r*th dtf-augmentation of *G* and let us create a digraph 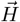 by orienting every edge *uv* ∈ *H* as follows:

1. If of *uv*, 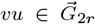, then orient *uv* in 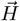 according to the corresponding arc in 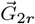 (if both arcs exists choose an arbitrary orientation),
2. otherwise there exists 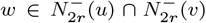 with *ω*_2*r*_ (*u*) + *ω*_2*r*_ (*v*) = dist_*G*_(*u,v*) ≤ 2*r*. Orient the edge *uv* towards that vertex *x* ∈ {*u,v*} for which *ω*_2*r*_(*x*) is larger.

We now argue that 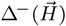 is small. Consider any vertex 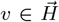. Every in-arc 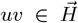 either is of type 1, of which we have at most 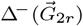, or of type 2. Consider a group of in-arcs *u_i_v*, 1 ≤ *i* ≤ *ℓ* of type 2 that are all present because of a common vertex *w*. Since 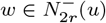, we have at most 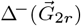 such groups. By construction, *ω*_2*r*_(*wu_i_*) ≤ *ω*_2*r*_ (*wv*) and since both weights sum to less than *2r*, this means that *ω*_2*r*_(*wu_i_*) ≤ *r*. Lemma 3 now tells us that *ℓ* ≤ *τ* +1. Therefore *v* has at most 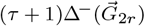 in-arcs of type 2 and we conclude that

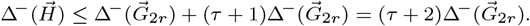

This finally implies that *H* is 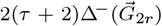-degenerate and therefore contains an independent set *A* ⊆ *V*(*H*) of size at least 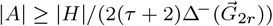. Taken together with the fact that 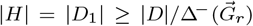 (every vertex added to *D*_1_ will cause at most 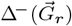 many vertices to be added to *D*_2_ in the loop at line 11 and *D* = *D*_1_ ∪ *D*_2_), we find that

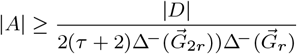

By construction of *H* we conclude that *A* is (2*r* + 1)-scattered in *G* of the claimed size.

Since a (2*r* + 1)-scattered set provides a lower bound for an *r*-dominating set, we conclude that Algorithm 1 computes a 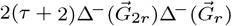-approximation of an optimal *r*-dominating set. In other words, we obtain a constant-factor approximation in graphs of bounded expansion since the quantities 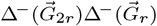 are constants per Theorem 1 in bounded expansion classes and the quantity 2(*τ* + 2) is a constant of our choosing.

In practice one could, depending on the value of 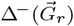 and 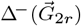, compute the optimal value for *τ* to minimize the approximation guarantee. However, this would necessitate the computation of 2*r* augmentation, the expensive step we want to avoid. Alternatively, we can choose a ‘good enough’ value for *τ* that still guarantees a constant-factor approximation while being easy to determine in practice. In the context of cDBGs, we found that *τ*:= (2*r*)^2^ yields reliably good results.

#### Computational Runtimes

See “Benchmarking” in Materials and Methods for benchmarking methods.

The `podarV` data set was retrieved from the NCBI SRA using accession SRR606249. The full build and indexing of the 103 million error-trimmed reads (10.3 Gbp in total) took approximately 23 minutes and required 12.8 GB of RAM. Loading the indices for search required 4.3 GB of RAM and a search with a 3 Mbp genome took approximately 32 seconds.

The `HuSBl` data set was retrieved from the NCBI SRA using accession SRR1976948. The full build and indexing of the 34 million error-trimmed reads (8.5 Gbp in total) required approximately 217 minutes and required 24.4 GB of RAM. Loading the indices for search required 18 GB of RAM and a search with a 3 Mbp genome took approximately 80 seconds.

For data set complexity (number of k-mers, number of cDBG nodes) please see Table 1.

#### spacegraphcats pipeline overview

spacegraphcats follows a series of steps when run on sequencing data, see Figure 5. In detail, we perform the following steps.

**Fig. 5.**
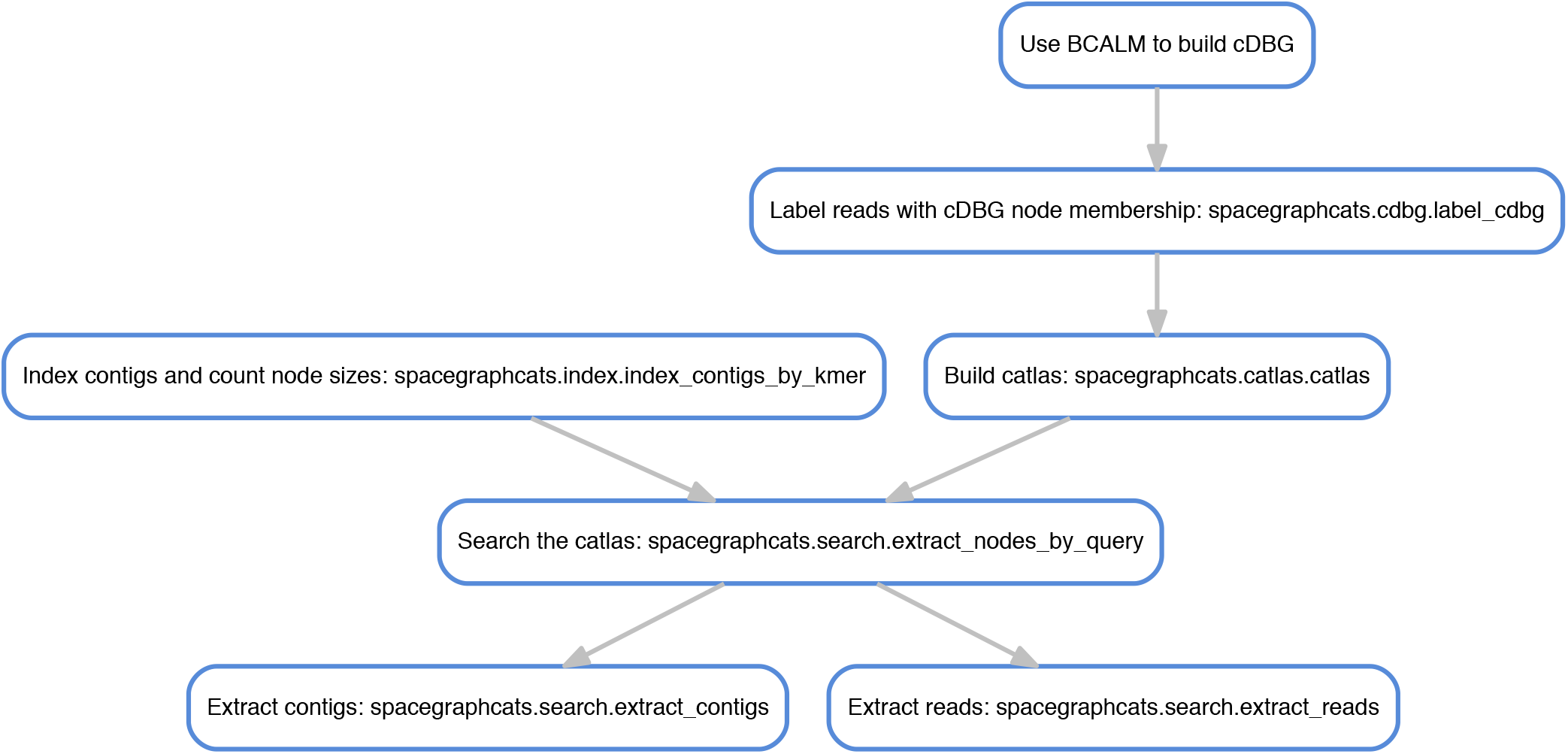
The steps followed by spacegraphcats when run on sequencing data.

*BCALM*. Use BCALM to generate a cDBG. Then convert a BCALM `unitigs.fa` output (a cDBG) into spacegraphcats files. Outputs an undirected graph, a file containing the sequences, and a `.info.csv` file containing information about the contig. Also outputs sourmash `k=31, scaled=1000` signatures for both input and output files.

*spacegraphcats.cdbg.label cdbg*. Build an index that can be used to retrieve individual reads or contigs by cDBG node ID; produce a SQLite database for fast retrieval. Briefly, this script creates a sqlite database with a single table, `sequences`, where a query like `SELECT DISTINCT sequences.offset FROM sequences WHERE label` … can be executed to return the offset of all sequences with the given label; the offsets refer to BGZF coordinates in the gzipped sequence collection. Here, ‘label’ is the cDBG ID to which the sequence belongs.

The script `extract_reads_by_frontier_sqlite.py` is a downstream script to extract the reads with a frontier search. Specifically: 1. walk through the contigs assembled from the cDBG; 2. build a DBG cover using khmer tags, such that every k-mer in the DBG is within distance d=40 of a tag; 3. label each tag with the cDBG node ID from the contig; 4. save for later use.

*spacegraphcats.catlas.catlas*. The catlas is a hierarchical atlas for querying graphs. Implements algorithms 1 and 2 (see main text).

*spacegraphcats.index.index_contigs_by_kmer*. Use Minimal Perfect Hashing (BBHash, https://github.com/rizkg/BBHash) to construct a fast lookup table connecting k-mers in the cDBG to cDBG node IDs. (BBHash reference: A. Limasset, G. Rizk, R. Chikhi, P. Peterlongo, Fast and Scalable Minimal Perfect Hashing for Massive Key Sets, SEA 2017.)

*spacegraphcats.search.extract_nodes_by_query*. Do a frontier search, and retrieve cDBG node IDs and MinHash signature for the retrieved contigs.

*spacegraphcats.search.extract_contigs*. Retrieve the unitig sequences for a given list of cDBG nodes. Consumes the output of `extract_nodes_by_query` to get the list of nodes.

*spacegraphcats.search.extract_reads*. Retrieve the reads for a list of cDBG nodes. Consumes the output of `extract_nodes_by_query` to get the list of nodes, and then uses the labeled cDBG output by `.cdbg.label_cdbg` to find reads that overlap with the unitigs in those nodes.

#### Query genome accession numbers for *Proteiniclasticum* search

See Table 3.

**Table 3.** Accession numbers for genomes used in *Proteiniclasticum* neighborhood query.

| Name | NCBI accession |
| --- | --- |
| *P. ruminis* CGMCC | GCA_900099635.1 |
| *P. ruminis* DSM | GCA_000701905.1 |
| *P. ruminis* ML2 | GCA_900115135.1 |

#### Amino Acid Identity results for *Proteiniclasticum*

See Table 4.

**Table 4.** CompareM results for *Proteiniclasticum* genomes. *P. ruminis shakya* is the result of assembling the reads extracted from `podarV` with the neighborhood search.

| Genome A | Genome B | Orthologous Genes | Mean AAI |
| --- | --- | --- | --- |
| *P. ruminis* ML2 | *P. ruminis* shakya | 2546 | 95.74 |
| *P. ruminis* DSM | *P. ruminis* shakya | 2391 | 93.47 |

#### HuSB1 analysis pipeline overview

See Figure 6. We implemented three workflows to analyze the plass-assembled `HuSB1` query neighborhoods.

**Fig. 6.**
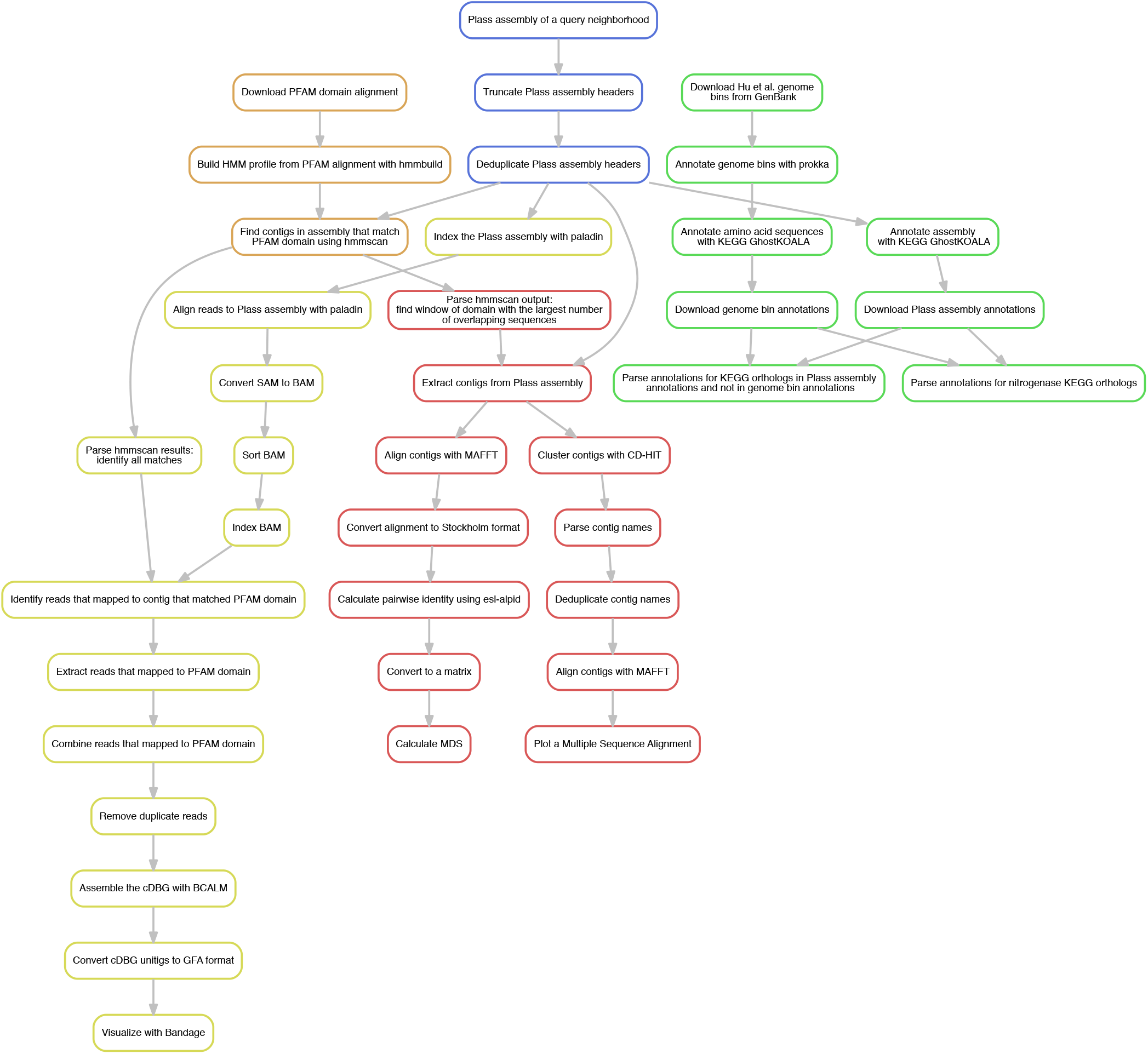
Three workflows implemented to analyze the plass-assembled HuSB1 query neighborhoods. The first three steps, depicted in blue, were common across all workflows. The green boxes depict the KEGG GhostKOALA annotation workflow, the results of which can be see in Figure 4. The orange boxes depict steps in common between the clustering and variant workflows used to generate Figure 4. The red boxes depict steps used to generate the MDS clustering plot and the multiple sequence alignment (see Figure 7). The gold boxes depict the steps of the variant workflow used to generate the assembly graphs.

#### Genome bin completeness improvements for HuSB1

See Table 5 and Table 6. Table 5 shows completeness metrics for the binned genomes from the `HuSB1` sample and used as queries. Table 6 shows completeness metrics for Plass assembled query neighborhoods after stringent read trimming at low abundance k-mers (k-mers present fewer than 5 times were removed) of the SB1 sample reads.

**Table 5.** Completeness metrics for the `HuSBl` genome bins from (35), used as queries into the SB1 metagenome. Completeness, redundancy, and strain heterogeneity as estimated by checkM, unique KEGG orthologs predicted by GhostKOALA, bin size in bases, and number of prokka-predicted protein sequences in the `HuSBl` bins. Table is ordered by completeness. Note we refer to the checkM term “contamination” as “redundancy” as this better describes the calculated metric.

| Species | Completeness (%) | Redundancy (%) | Strain heterogeneity (%) | Unique KOs | Size (bases) | Number of proteins |
| --- | --- | --- | --- | --- | --- | --- |
| WS6 bacterium 36_33 | 31.5 | 6.5 | 85.7 | 127 | 439774 | 423 |
| P. bacterium 34_609 | 34.0 | 1.1 | 12.5 | 173 | 567617 | 557 |
| P. bacterium 33_209 | 47.9 | 5.2 | 0 | 140 | 510490 | 526 |
| M. bacterium 39_7 | 50.1 | 5.9 | 0 | 208 | 708389 | 657 |
| P. acetatigenes isolate 50_10 | 56.7 | 3.6 | 66.7 | 506 | 1716233 | 1510 |
| WS6 bacterium 34_10 | 61.9 | 29.5 | 61.3 | 219 | 1127556 | 1065 |
| M. infera isolate 46_47 | 63.8 | 0.9 | 100 | 502 | 1225111 | 1170 |
| A. bacterium 34_128 | 64.4 | 5.1 | 0 | 423 | 894916 | 785 |
| A. thermophila isolate 46_16 | 67.2 | 16.4 | 0 | 474 | 1524726 | 1350 |
| A. bacterium 49_20 | 69.6 | 8.3 | 0 | 380 | 1023183 | 891 |
| M. marisnigri isolate 62_101 | 72.1 | 1.0 | 0 | 583 | 1592820 | 1742 |
| M. bacterium 46_47 | 72.9 | 0.1 | 0 | 563 | 1629409 | 1413 |
| B. bacterium 38_7 | 80.0 | 2.7 | 0 | 601 | 2137321 | 1697 |
| Methanocalculus sp. 52_23 | 82.7 | 4.6 | 66.7 | 689 | 1973787 | 2074 |
| Desulfotomaculum sp. 46_80 | 83.5 | 2.5 | 42.9 | 834 | 2251381 | 2148 |
| S. bacterium 57_84 | 90.8 | 1.9 | 50 | 608 | 1255134 | 1277 |
| S. bacterium 53_16 | 91.5 | 4.2 | 0 | 746 | 1772227 | 1741 |
| Desulfotomaculum sp. 46_296 | 91.5 | 9.0 | 65.4 | 851 | 2328136 | 2265 |
| A. bacterium 66_15 | 94.2 | 2.8 | 20 | 811 | 2228088 | 2185 |
| C. bacterium 38_11 | 94.4 | 0.3 | 0 | 825 | 1882878 | 1744 |
| TA06 bacterium 32_111 | 94.5 | 0 | 0 | 619 | 1861827 | 1736 |
| Methanobacterium sp. 42_16 | 97.6 | 0 | 0 | 769 | 2173293 | 2149 |
| M. harundinacea isolate 57_489 | 100 | 0 | 0 | 814 | 2382964 | 2377 |

**Table 6.** Completeness metrics for the query neighborhoods extracted from the `HuSB1` metagenome by spacegraphcats. Completeness, redundancy, and strain heterogeneity as estimated by checkM, unique KEGG orthologs predicted by GhostKOALA, neighborhood size in bases, and number of plass-assembled protein sequences in the query neighborhoods. Table is ordered by completeness of query bins (see Table 5). All estimates were performed on the k-mer trimmed (k >= 5)Plass-assembled proteins except size of neighborhood in bases, for which we used the neighborhood unitig sequences output by spacegraphcats.

| Species | Completeness (%) | Redundancy (%) | Strain heterogeneity (%) | Unique KOs | Size (bases) | Number of proteins |
| --- | --- | --- | --- | --- | --- | --- |
| WS6 bacterium 36_33 | 45.7 | 1675.8 | 87.9 | 140 | 1206915 | 12990 |
| P. bacterium 34_609 | 39.7 | 637.3 | 74.8 | 239 | 3604616 | 12372 |
| P. bacterium 33_209 | 46.0 | 516.1 | 92.7 | 160 | 1830403 | 9836 |
| M. bacterium 39_7 | 30.8 | 1.9 | 20 | 121 | 1356859 | 1208 |
| P. acetatigenes isolate 50_10 | 42.7 | 66.3 | 86.7 | 557 | 6829683 | 6135 |
| WS6 bacterium 34_10 | 58.6 | 352.8 | 86.0 | 219 | 2840498 | 6549 |
| M. infera isolate 46_47 | 67.0 | 309.8 | 92.6 | 592 | 4021633 | 11798 |
| A. bacterium 34_128 | 65.5 | 332 | 58.9 | 446 | 2703261 | 9030 |
| A. thermophila isolate 46_16 | 60.5 | 250 | 88.5 | 457 | 5080416 | 13780 |
| A. bacterium 49_20 | 71.3 | 187.2 | 76.7 | 392 | 4166615 | 6040 |
| M. marisnigri isolate 62_101 | 82.2 | 1067.9 | 89.9 | 687 | 11380474 | 49119 |
| M. bacterium 46_47 | 81.0 | 251.3 | 97.5 | 629 | 5308863 | 15964 |
| B. bacterium 38_7 | 44.0 | 5.3 | 20 | 508 | 4229976 | 3901 |
| Methanocalculus sp. 52_23 | 89.3 | 443.4 | 87.0 | 741 | 8018167 | 20999 |
| Desulfotomaculum sp. 46_80 | 94.8 | 2327.3 | 94.7 | 858 | 7969109 | 48841 |
| S. bacterium 57_84 | 90.8 | 598.2 | 83.9 | 686 | 5968936 | 24774 |
| S. bacterium 53_16 | 94.0 | 95.6 | 83.9 | 786 | 7145072 | 12138 |
| Desulfotomaculum sp. 46_296 | 94.8 | 2535.0 | 80.4 | 888 | 17836454 | 48174 |
| A. bacterium 66_15 | 83.6 | 10.6 | 83.7 | 792 | 10865439 | 8152 |
| C. bacterium 38_11 | 86.7 | 18.2 | 78.4 | 800 | 4932373 | 4582 |
| TA06 bacterium 32_111 | 95.6 | 45.7 | 88.2 | 626 | 3811703 | 5724 |
| Methanobacterium sp. 42_16 | 92.2 | 31.3 | 90.9 | 782 | 7007709 | 6335 |
| M. harundinacea isolate 57_489 | 98.8 | 355.6 | 89.6 | 841 | 35722445 | 35093 |

#### K-mer inclusion of reads by MEGAHIT assemblies

See Table 8. We estimated the number of k-mers in each query neighborhood that were contained in the MEGAHIT assembly of that query neighborhood. We used sourmash `compute` to calculate signatures with k-size of 31 and a scaled value of 2000. We then used sourmash `compare` to estimate containment in MEGAHIT assemblies. The query neighborhood with the smallest containment, *M. harundi-nacea* isolate *57_489*, had the largest query neighborhood, while the query neighborhood with the largest containment, *M. bacterium 39_7*, had the smallest query neighborhood.

**Table 7.**
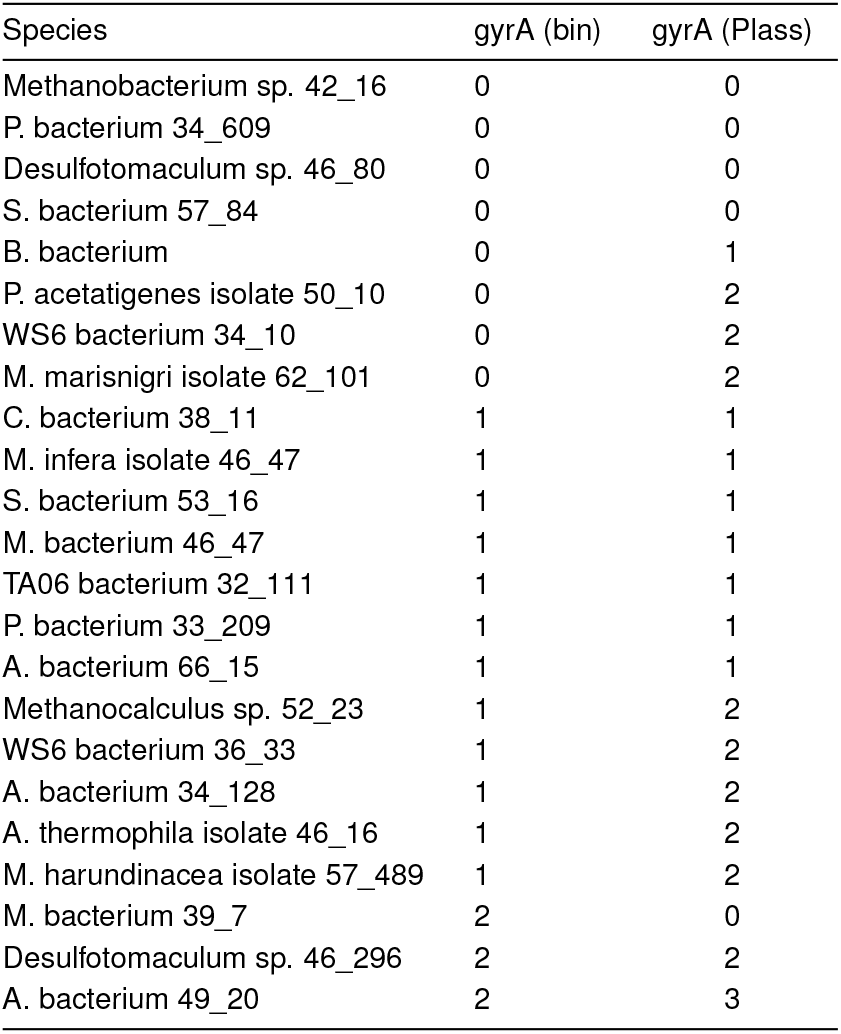
Bin and neighborhood gyrA protein content. gyrA count for each bin is the number of gyrA amino acid sequences that are part of the original bin. gyrA count by Plass is the minimum number of gyrA amino acid sequences supported by at least one position with at least 10 copies of each variant, e.g. “3” indicates that there is at least one position in the multiple sequence alignment of gyrA sequences for that neighborhood that has 3 distinct variants in 10 distinct sequences.

**Table 8.** Containment of neighborhood k-mer content in MEGAHIT nucleotide assemblies.

| Species | MEGAHIT assembly containment |
| --- | --- |
| M. harundinacea isolate 57_489 | 4.2% |
| Desulfotomaculum sp. 46_296 | 12.7% |
| M. marisnigri isolate 62_101 | 13.6% |
| S. bacterium 57_84 | 19.4% |
| P. bacterium 34_609 | 19.7% |
| A. bacterium 66_15 | 20.5% |
| Desulfotomaculum sp. 46_80 | 24.1% |
| P. bacterium 33_209 | 26.3% |
| S. bacterium 53_16 | 30.9% |
| A. bacterium 49_20 | 31.9% |
| Methanocalculus sp. 52_23 | 33.4% |
| M. bacterium 46_47 | 36.6% |
| P. acetatigenes isolate 50_10 | 36.6% |
| A. bacterium 34_128 | 36.8% |
| M. infera isolate 46_47 | 38.0% |
| Methanobacterium sp. 42_16 | 38.0% |
| A. thermophila isolate 46_16 | 38.6% |
| TA06 bacterium 32_111 | 44.1% |
| C. bacterium 38_11 | 44.4% |
| WS6 bacterium 34_10 | 53.2% |
| WS6 bacterium 36_33 | 53.8% |
| B. bacterium | 54.2% |
| M. bacterium 39_7 | 55.7% |

#### gyrA alignment

See Figure 7. The MDS plot in the left panel of figure 4 shows distinct gyrA sequences identified in the Plass assemblies using HMMER. To visualize the sequences within these clusters and in other query neighborhoods, we constructed a multiple sequence alignment. However, because many sequences assembled by Plass were fragmented (see Results: Some query neighborhoods contain substantial strain variation), we first clustered the sequences at 95% similarity using CD-HIT. We then aligned the centroid sequences using MAFFT with default settings. To produce the multiple sequence alignment visualization, we calculated an unrooted neighbor joining tree using the MAFFT alignment. Then we used the function msaplot in the R package ggtree to plot the alignment.

**Fig. 7.**
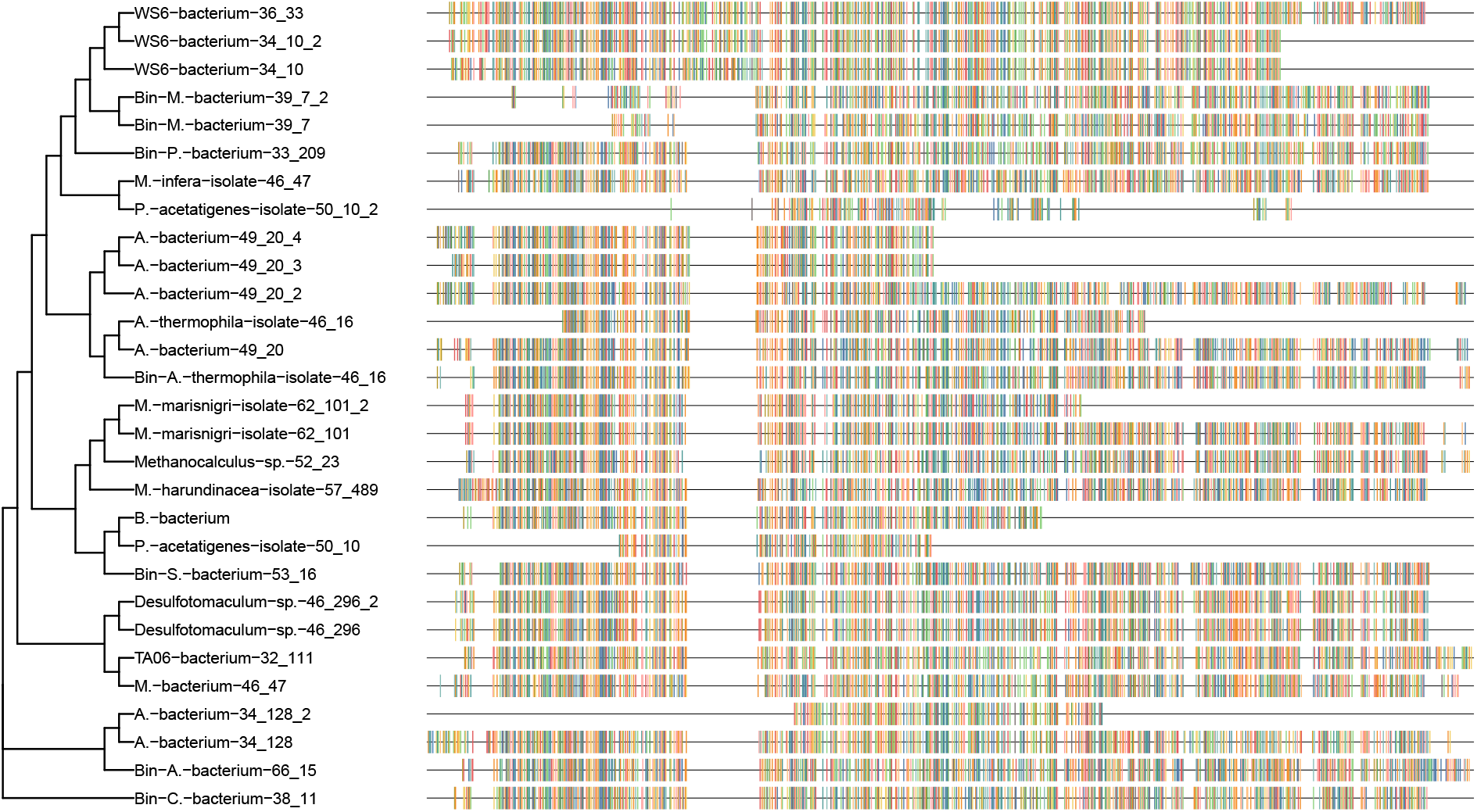
A multiple sequence alignment and neighbor joining tree of representative gyrA amino acid fragments assembled by Plass from the genome neighborhoods in HuSB1. Protein sequences that originated from the genome bin are prepended with “Bin.” All other sequences were assembled by Plass.

#### gyrA by neighborhood

See Table 7. As can be seen in the left panel of Figure 4 in the main text, we observe many unique amino acid sequences per single copy ortholog per query neighborhood. Although we observe many possible traversal paths in compact De Bruijn graphs built from reads that give rise to these sequences, we have no way to ascertain whether we observed combinatorial complexity by assembling variants that would never be linked in nature. Therefore, we sought to conservatively estimate the number of positions per amino acid sequence that contained variants using MAFFT alignments. First, we subsetted the alignment to sequences from one query neighborhood. Then we identified all aligned nongap characters for each position in the alignment (gaps were induced in some neighborhoods by the presence of amino acid residues in other query neighborhood amino acid sequences). For each of these positions, we counted the number of unique amino acid sequences per position, and the number of times each occurred at that position. We then elimated any variant that occurred fewer than 10 times. Lastly, we counted the number of well-supported distinct characters. We did this for gyrA, as well as the amino acid sequences for the other genes we tested (see other genes). Table 7 shows that we see increased number of gyrA sequences in many neighborhoods even with this conservative approach.

#### Other genes

See bin and neighborhood content results for *alaS* in Table 10, *gyrB* in Table 11, *recA* in Table 12, *rpb2d6* in Table 13, rplB in Table 14, and rpsC in Table 15. We selected *gyrB* and *recA* because they were used by HuSB1 to assign taxonomy to binned genomes. We selected other genes used as single copy orthologs by programs like CheckM, and with longer PFAM domains.

**Table 9.** Protein names and PFAM accessions for targeted analyses.

| Name | PFAM accession |
| --- | --- |
| recA | PF00154 |
| rplB | PF00181 |
| rpsC | PF00189 |
| gyrB | PF00204 |
| gyrA | PF00521 |
| rpb2d6 | PF00562 |
| alaS | PF01411 |

**Table 10.** Bin and neighborhood alaS protein content.

| Species | alaS (bin) | alaS (Plass) |
| --- | --- | --- |
| P. acetatigenes isolate 50_10 | 0 | 0 |
| A. bacterium 49_20 | 0 | 0 |
| P. bacterium 34_609 | 0 | 0 |
| B. bacterium | 0 | 0 |
| S. bacterium 53_16 | 0 | 0 |
| A. bacterium 34_128 | 0 | 0 |
| M. infera isolate 46_47 | 0 | 2 |
| M. marisnigri isolate 62_101 | 0 | 2 |
| M. bacterium 39_7 | 1 | 0 |
| Methanobacterium sp. 42_16 | 1 | 1 |
| C. bacterium 38_11 | 1 | 1 |
| S. bacterium 57_84 | 1 | 1 |
| TA06 bacterium 32_111 | 1 | 1 |
| P. bacterium 33_209 | 1 | 1 |
| A. bacterium 66_15 | 1 | 1 |
| M. harundinacea isolate 57_489 | 1 | 1 |
| Methanocalculus sp. 52_23 | 1 | 2 |
| WS6 bacterium 36_33 | 1 | 2 |
| Desulfotomaculum sp. 46_80 | 1 | 2 |
| M. bacterium 46_47 | 1 | 2 |
| Desulfotomaculum sp. 46_296 | 1 | 2 |
| A. thermophila isolate 46_16 | 2 | 1 |
| WS6 bacterium 34_10 | 2 | 2 |

**Table 11.** Bin and neighborhood gyrB protein content.

| Species | gyrB (bin) | gyrB (Plass) |
| --- | --- | --- |
| M. bacterium 39_7 | 0 | 0 |
| P. acetatigenes isolate 50_10 | 0 | 0 |
| Methanobacterium sp. 42_16 | 0 | 0 |
| WS6 bacterium 36_33 | 0 | 0 |
| P. bacterium 34_609 | 0 | 0 |
| Desulfotomaculum sp. 46_80 | 0 | 0 |
| S. bacterium 57_84 | 0 | 0 |
| S. bacterium 53_16 | 0 | 0 |
| A. thermophila isolate 46_16 | 0 | 2 |
| P. bacterium 33_209 | 1 | 0 |
| C. bacterium 38_11 | 1 | 1 |
| M. infera isolate 46_47 | 1 | 1 |
| M. bacterium 46_47 | 1 | 1 |
| TA06 bacterium 32_111 | 1 | 1 |
| A. bacterium 66_15 | 1 | 1 |
| M. harundinacea isolate 57_489 | 1 | 1 |
| WS6 bacterium 34_10 | 1 | 2 |
| Methanocalculus sp. 52_23 | 1 | 2 |
| M. marisnigri isolate 62_101 | 1 | 2 |
| A. bacterium 34_128 | 1 | 2 |
| A. bacterium 49_20 | 2 | 2 |
| B. bacterium | 2 | 2 |
| Desulfotomaculum sp. 46_296 | 2 | 2 |

**Table 12.** Bin and neighborhood recA protein content.

| Species | recA (bin) | recA (Plass) |
| --- | --- | --- |
| M. bacterium 39_7 | 0 | 0 |
| WS6 bacterium 34_10 | 0 | 0 |
| Methanocalculus sp. 52_23 | 0 | 0 |
| A. bacterium 49_20 | 0 | 0 |
| WS6 bacterium 36_33 | 0 | 0 |
| P. bacterium 34_609 | 0 | 0 |
| M. marisnigri isolate 62_101 | 0 | 0 |
| S. bacterium 53_16 | 0 | 0 |
| M. bacterium 46_47 | 0 | 0 |
| A. thermophila isolate 46_16 | 0 | 1 |
| B. bacterium | 1 | 0 |
| P. acetatigenes isolate 50_10 | 1 | 1 |
| Methanobacterium sp. 42_16 | 1 | 1 |
| C. bacterium 38_11 | 1 | 1 |
| M. infera isolate 46_47 | 1 | 1 |
| S. bacterium 57_84 | 1 | 1 |
| A. bacterium 34_128 | 1 | 1 |
| TA06 bacterium 32_111 | 1 | 1 |
| P. bacterium 33_209 | 1 | 1 |
| A. bacterium 66_15 | 1 | 1 |
| M. harundinacea isolate 57_489 | 1 | 1 |
| Desulfotomaculum sp. 46_80 | 1 | 2 |
| Desulfotomaculum sp. 46_296 | 1 | 2 |

**Table 13.** Bin and neighborhood rpb2d6 protein content.

| Species | rpb2d6 (bin) | rpb2d6 (Plass) |
| --- | --- | --- |
| P. acetatigenes isolate 50_10 | 0 | 0 |
| P. bacterium 34_609 | 0 | 0 |
| S. bacterium 57_84 | 0 | 1 |
| M. bacterium 46_47 | 0 | 1 |
| C. bacterium 38_11 | 1 | 0 |
| A. bacterium 49_20 | 1 | 0 |
| M. bacterium 39_7 | 1 | 1 |
| Methanobacterium sp. 42_16 | 1 | 1 |
| Methanocalculus sp. 52_23 | 1 | 1 |
| M. infera isolate 46_47 | 1 | 1 |
| B. bacterium | 1 | 1 |
| S. bacterium 53_16 | 1 | 1 |
| A. bacterium 34_128 | 1 | 1 |
| TA06 bacterium 32_111 | 1 | 1 |
| A. bacterium 66_15 | 1 | 1 |
| A. thermophila isolate 46_16 | 1 | 1 |
| WS6 bacterium 36_33 | 1 | 2 |
| Desulfotomaculum sp. 46_80 | 1 | 2 |
| M. marisnigri isolate 62_101 | 1 | 2 |
| Desulfotomaculum sp. 46_296 | 1 | 2 |
| P. bacterium 33_209 | 1 | 2 |
| M. harundinacea isolate 57_489 | 1 | 2 |
| WS6 bacterium 34_10 | 2 | 2 |

**Table 14.**
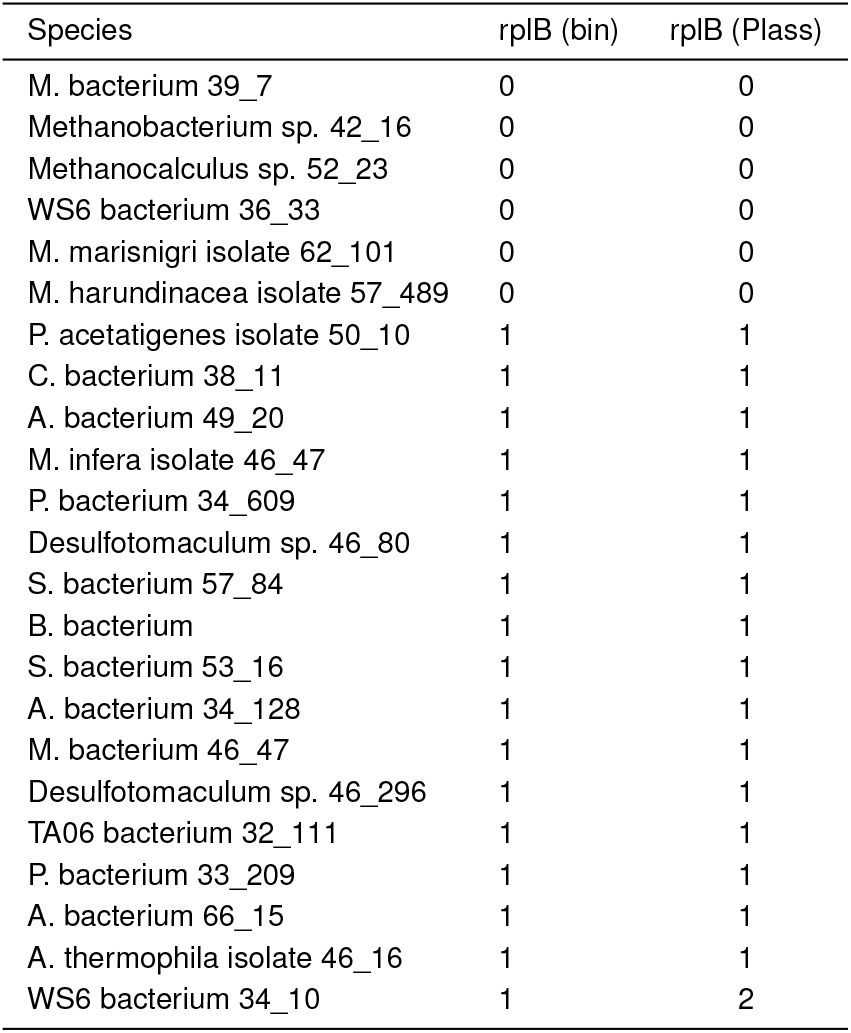
Bin and neighborhood rplB protein content.

| Species | rplB (bin) | rplB (Plass) |
| --- | --- | --- |
| M. bacterium 39_7 | 0 | 0 |
| Methanobacterium sp. 42_16 | 0 | 0 |
| Methanocalculus sp. 52_23 | 0 | 0 |
| WS6 bacterium 36_33 | 0 | 0 |
| M. marisnigri isolate 62_101 | 0 | 0 |
| M. harundinacea isolate 57_489 | 0 | 0 |
| P. acetatigenes isolate 50_10 | 1 | 1 |
| C. bacterium 38_11 | 1 | 1 |
| A. bacterium 49_20 | 1 | 1 |
| M. infera isolate 46_47 | 1 | 1 |
| P. bacterium 34_609 | 1 | 1 |
| Desulfotomaculum sp. 46_80 | 1 | 1 |
| S. bacterium 57_84 | 1 | 1 |
| B. bacterium | 1 | 1 |
| S. bacterium 53_16 | 1 | 1 |
| A. bacterium 34_128 | 1 | 1 |
| M. bacterium 46_47 | 1 | 1 |
| Desulfotomaculum sp. 46_296 | 1 | 1 |
| TA06 bacterium 32_111 | 1 | 1 |
| P. bacterium 33_209 | 1 | 1 |
| A. bacterium 66_15 | 1 | 1 |
| A. thermophila isolate 46_16 | 1 | 1 |
| WS6 bacterium 34_10 | 1 | 2 |

**Table 15.** Bin and neighborhood rpsC protein content.

| Species | rpsC (bin) | rpsC (Plass) |
| --- | --- | --- |
| M. bacterium 39_7 | 0 | 0 |
| P. acetatigenes isolate 50_10 | 0 | 0 |
| WS6 bacterium 34_10 | 0 | 0 |
| Methanobacterium sp. 42_16 | 0 | 0 |
| Methanocalculus sp. 52_23 | 0 | 0 |
| WS6 bacterium 36_33 | 0 | 0 |
| M. marisnigri isolate 62_101 | 0 | 0 |
| B. bacterium | 0 | 0 |
| P. bacterium 33_209 | 0 | 0 |
| M. harundinacea isolate 57_489 | 0 | 0 |
| M. infera isolate 46_47 | 0 | 1 |
| C. bacterium 38_11 | 1 | 1 |
| A. bacterium 49_20 | 1 | 1 |
| P. bacterium 34_609 | 1 | 1 |
| S. bacterium 57_84 | 1 | 1 |
| S. bacterium 53_16 | 1 | 1 |
| A. bacterium 34_128 | 1 | 1 |
| M. bacterium 46_47 | 1 | 1 |
| TA06 bacterium 32_111 | 1 | 1 |
| A. bacterium 66_15 | 1 | 1 |
| A. thermophila isolate 46_16 | 1 | 1 |
| Desulfotomaculum sp. 46_80 | 1 | 2 |
| Desulfotomaculum sp. 46_296 | 1 | 2 |

## Footnotes

* an invertible function that defines both an index and the corresponding inverted index

† That is, graphs about which we make no structural assumptions.

## References

1. Christopher Quince, Alan W Walker, Jared T Simpson, Nicholas J Loman, and Nicola Segata. Shotgun metagenomics, from sampling to analysis. Nature Biotechnology, 35(9):833–844, sep 2017.. URL https://doi.org/10.1038/nbt.3935.

2. Jason Pell et al. Scaling metagenome sequence assembly with probabilistic de bruijn graphs. PNAS, 109(33):13272–13277, 2012.. URL https://doi.org/10.1073/pnas.1121464109.

3. C. C. Laczny, C. Kiefer, V. Galata, T. Fehlmann, C. Backes, and A. Keller. Busybee web: metagenomic data analysis by bootstrapped supervised binning and annotation. Nucleic Acids Research, page gkx348, 2017.

4. H. Lin and Y. Liao. Accurate binning of metagenomic contigs via automated clustering sequences using information of genomic signatures and marker genes. Scientific Reports, 6: 24175, 2016.

5. Donovan H. Parks, Christian Rinke, Maria Chuvochina, Pierre-Alain Chaumeil, Ben J. Woodcroft, Paul N. Evans, Philip Hugenholtz, and Gene W. Tyson. Recovery of nearly 8, 000 metagenome-assembled genomes substantially expands the tree of life. Nature Microbiology, 2(11):1533–1542, sep 2017.. URL https://doi.org/10.1038/s41564-017-0012-7.

6. Benjamin J. Tully, Elaina D. Graham, and John F. Heidelberg. The reconstruction of 2, 631 draft metagenome-assembled genomes from the global oceans. Scientific Data, 5:170203, jan 2018.. URL https://doi.org/10.1038/sdata.2017.203.

7. Robert D. Stewart, Marc D. Auffret, Amanda Warr, Andrew H. Wiser, Maximilian O. Press, Kyle W. Langford, Ivan Liachko, Timothy J. Snelling, Richard J. Dewhurst, Alan W. Walker, Rainer Roehe, and Mick Watson. Assembly of 913 microbial genomes from metagenomic sequencing of the cow rumen. Nature Communications, 9(1), feb 2018.. URL https://doi.org/10.1038/s41467-018-03317-6.

8. Tom O. Delmont, Christopher Quince, Alon Shaiber, Özcan C. Esen, Sonny TM Lee, Michael S. Rappé, Sandra L. McLellan, Sebastian Lücker, and A. Murat Eren. Nitrogenfixing populations of planctomycetes and proteobacteria are abundant in surface ocean metagenomes. Nature Microbiology, 3(7):804–813, jun 2018.. URL https://doi.org/10.1038/s41564-018-0176-9.

9. Laura A. Hug, Brett J. Baker, Karthik Anantharaman, Christopher T. Brown, Alexander J. Probst, Cindy J. Castelle, Cristina N. Butterfield, Alex W. Hernsdorf, Yuki Amano, Kotaro Ise, Yohey Suzuki, Natasha Dudek, David A. Relman, Kari M. Finstad, Ronald Amundson, Brian C. Thomas, and Jillian F. Banfield. A new view of the tree of life. Nature Microbiology, 1(5), apr 2016.. URL https://doi.org/10.1038/nmicrobiol.2016.48.

10. Edoardo Pasolli, Francesco Asnicar, Serena Manara, Moreno Zolfo, Nicolai Karcher, Federica Armanini, Francesco Beghini, Paolo Manghi, Adrian Tett, Paolo Ghensi, Maria Carmen Collado, Benjamin L. Rice, Casey DuLong, Xochitl C. Morgan, Christopher D. Golden, Christopher Quince, Curtis Huttenhower, and Nicola Segata. Extensive unexplored human microbiome diversity revealed by over 150, 000 genomes from metagenomes spanning age, geography, and lifestyle. Cell, 176(3):649–662.e20, jan 2019.. URL https://doi.org/10.1016/j.cell.2019.01.001.

11. Alexander Sczyrba, Peter Hofmann, Peter Belmann, David Koslicki, Stefan Janssen, Johannes Dröge, Ivan Gregor, Stephan Majda, Jessika Fiedler, Eik Dahms, Andreas Bremges, Adrian Fritz, Ruben Garrido-Oter, Tue Sparholt Jørgensen, Nicole Shapiro, Philip D Blood, Alexey Gurevich, Yang Bai, Dmitrij Turaev, Matthew Z DeMaere, Rayan Chikhi, Niranjan Na-garajan, Christopher Quince, Fernando Meyer, Monika Balvočiūtė, Lars Hestbjerg Hansen, Søren J Sørensen, Burton K H Chia, Bertrand Denis, Jeff L Froula, Zhong Wang, Robert Egan, Dongwan Don Kang, Jeffrey J Cook, Charles Deltel, Michael Beckstette, Claire Lemaitre, Pierre Peterlongo, Guillaume Rizk, Dominique Lavenier, Yu-Wei Wu, Steven W Singer, Chirag Jain, Marc Strous, Heiner Klingenberg, Peter Meinicke, Michael D Barton, Thomas Lingner, Hsin-Hung Lin, Yu-Chieh Liao, Genivaldo Gueiros Z Silva, Daniel A Cuevas, Robert A Edwards, Surya Saha, Vitor C Piro, Bernhard Y Renard, Mihai Pop, Hans-Peter Klenk, Markus Göker, Nikos C Kyrpides, Tanja Woyke, Julia A Vorholt, Paul Schulze-Lefert, Edward M Rubin, Aaron E Darling, Thomas Rattei, and Alice C McHardy. Critical assessment of metagenome interpretation—a benchmark of metagenomics software. Nature Methods, 14 (11):1063–1071, oct 2017.. URL https://doi.org/10.1038/nmeth.4458.

12. Sherine Awad, Luiz Irber, and C. Titus Brown. Evaluating metagenome assembly on a simple defined community with many strain variants. https://www.biorxiv.org/content/early/2017/07/03/155358, 2017. URL https://www.biorxiv.org/content/early/2017/07/03/155358.

13. C Titus Brown. Strain recovery from metagenomes. Nature Biotechnology, 33(10):1041–1043, oct 2015.. URL https://doi.org/10.1038/nbt.3375.

14. Ilana L. Brito and Eric J. Alm. Tracking strains in the microbiome: Insights from metagenomics and models. Frontiersin Microbiology, 7, may 2016.. URL https://doi.org/10.3389/fmicb.2016.00712.

15. Johannes Alneberg, Christofer M. G. Karlsson, Anna-Maria Divne, Claudia Bergin, Felix Homa, Markus V. Lindh, Luisa W. Hugerth, Thijs J. G. Ettema, Stefan Bertilsson, Anders F. Andersson, and Jarone Pinhassi. Genomes from uncultivated prokaryotes: a comparison of metagenome-assembled and single-amplified genomes. Microbiome, 6(1), sep 2018.. URL https://doi.org/10.1186/s40168-018-0550-0.

16. Christopher Quince, Tom O. Delmont, Sébastien Raguideau, Johannes Alneberg, Aaron E. Darling, Gavin Collins, and A. Murat Eren. DESMAN: a new tool for de novo extraction of strains from metagenomes. Genome Biology, 18(1), sep 2017.. URL https://doi.org/10.1186/s13059-017-1309-9.

17. Stephen Nayfach, Beltran Rodriguez-Mueller, Nandita Garud, and Katherine S. Pollard. An integrated metagenomics pipeline for strain profiling reveals novel patterns of bacterial transmission and biogeography. Genome Research, 26(11):1612–1625, oct 2016.. URL https://doi.org/10.1101/gr.201863.115.

18. Erik Garrison. Graphical pangenomics. PhD thesis, Cambridge University, October 2018. URL https://doi.org/10.5281/zenodo.1463032. As submitted, awaiting viva (defense) and further revision.

19. Florian Plaza Onate, Emmanuelle Le Chatelier, Mathieu Almeida, Alessandra C L Cervino, Franck Gauthier, Frederic Magoules, S Dusko Ehrlich, and Matthieu Pichaud. MSPminer: abundance-based reconstitution of microbial pan-genomes from shotgun metagenomic data. Bioinformatics, sep 2018.. URL https://doi.org/10.1093/bioinformatics/bty830.

20. Jillian M. Petersen, Anna Kemper, Harald Gruber-Vodicka, Ulisse Cardini, Matthijs van der Geest, Manuel Kleiner, Silvia Bulgheresi, Marc Mußmann, Craig Herbold, Brandon K.B. Seah, Chakkiath Paul Antony, Dan Liu, Alexandra Belitz, and Miriam Weber. Chemosynthetic symbionts of marine invertebrate animals are capable of nitrogen fixation. Nature Microbiology, 2(1), oct 2016.. URL https://doi.org/10.1038/nmicrobiol.2016.195.

21. Evgenii I Olekhnovich, Artem T Vasilyev, Vladimir I Ulyantsev, Elena S Kostryukova, and Alexander V Tyakht. MetaCherchant: analyzing genomic context of antibiotic resistance genes in gut microbiota. Bioinformatics, 34(3):434–444, oct 2017.. URL https://doi.org/10.1093/bioinformatics/btx681.

22. Tyler P. Barnum, Israel A. Figueroa, Charlotte I. Carlström, Lauren N. Lucas, Anna L. Engelbrektson, and John D. Coates. Genome-resolved metagenomics identifies genetic mobility, metabolic interactions, and unexpected diversity in perchlorate-reducing communities. The ISME Journal, 12(6):1568–1581, feb 2018.. URL https://doi.org/10.1038/s41396-018-0081-5.

23. C. T. Brown, D. Moritz, M. P. O’Brien, F. Reidl, and B. D. Sullivan. spacegraphcats, v1.0. http://dx.doi.org/10.5281/zenodo.1478025, November 2018.

24. F. Reidl. Structural sparseness and complex networks. 2016. URL http://publications.rwth-aachen.de/record/565064. Aachen, Techn. Hochsch., Diss., 2015.

25. Richard M Karp. Reducibility among combinatorial problems. In Complexity of computer computations, pages 85–103. Springer, 1972.

26. Miroslav Chlebík and Janka Chlebíková. Approximation hardness of dominating set problems in bounded degree graphs. Information and Computation, 206(11):1264–1275, 2008.

27. Rodney G Downey and Michael Ralph Fellows. Parameterized complexity. Springer Science & Business Media, 2012.

28. Patrice Ossona de Mendez et al. Sparsity: graphs, structures, and algorithms, volume 28. Springer Science & Business Media, 2012..

29. Antoine Limasset, Guillaume Rizk, Rayan Chikhi, and Pierre Peterlongo. Fast and scalable minimal perfect hashing for massive key sets. CoRR, abs/1702.03154, 2017. URL http://arxiv.org/abs/1702.03154.

30. M. Shakya, C. Quince, J. H. Campbell, Z. K. Yang, C. W. Schadt, and M. Podar. Comparative metagenomic and rrna microbial diversity characterization using archaeal and bacterial synthetic communities. Environmental Microbiology, 15(6):1882–1899, 2013. ISSN 1462-2920.. URL http://dx.doi.org/10.1111/1462-2920.12086.

31. Dinghua Li, Ruibang Luo, Chi-Man Liu, Chi-Ming Leung, Hĩng Fung Ting, Kunihiko Sadakane, Hiroshi Yamashita, and Tak-Wah Lam. MEGAHIT v1.0: A fast and scalable metagenome assembler driven by a dvanced methodologies and community practices. Methods, 102:3–11, jun 2016.. URL https://doi.org/10.1016/j.ymeth.2016.02.020.

32. Brandon K. B. Seah and Harald R. Gruber-Vodicka. gbtools: Interactive visualization of metagenome bins in r. Frontiers in Microbiology, 6, dec 2015.. URL https://doi.org/10.3389/fmicb.2015.01451.

33. Sergey Nurk, Dmitry Meleshko, Anton Korobeynikov, and Pavel A Pevzner. metaspades: a new versatile metagenomic assembler. Genome Research, 27(5):824–834, 2017.

34. Itai Sharon, Michael Kertesz, Laura A. Hug, Dmitry Pushkarev, Timothy A. Blauwkamp, Cindy J. Castelle, Mojgan Amirebrahimi, Brian C. Thomas, David Burstein, Susannah G. Tringe, Kenneth H. Williams, and Jillian F. Banfield. Accurate, multi-kb reads resolve complex populations and detect rare microorganisms. Genome Research, 24(4):534–543, feb 2015.. URL https://doi.org/10.1101/gr.183012.114.

35. Ping Hu, Lauren Tom, Andrea Singh, Brian C. Thomas, Brett J. Baker, Yvette M. Piceno, Gary L. Andersen, and Jillian F. Banfield. Genome-resolved metagenomic analysis reveals roles for candidate phyla and other microbial community members in biogeochemical transformations in oil reservoirs. mBio, 7(1), jan 2016.. URL https://doi.org/10.1128/mbio.01669-15.

36. Martin Steinegger, Milot Mirdita, and Johannes Soding. Protein-level assembly increases protein sequence recovery from metagenomic samples manyfold. aug 2018.. URL https://doi.org/10.1101/386110.

37. Youngik Yang and Shibu Yooseph. SPA: a short peptide assembler for metagenomic data. Nucleic Acids Research, 41(8):e91–e91, feb 2013.. URL https://doi.org/10.1093/nar/gkt118.

38. Donovan H. Parks, Michael Imelfort, Connor T. Skennerton, Philip Hugenholtz, and Gene W. Tyson. CheckM: assessing the quality of microbial genomes recovered from isolates, single cells, and metagenomes. Genome Research, 25(7):1043–1055, may 2015.. URL https://doi.org/10.1101/gr.186072.114.

39. P. Hu, L. Tom, A. Singh, B. C. Thomas, B. J. Baker, Y. M. Piceno, G. L. Andersen, and J.F. Banfield. Genome-resolved metagenomic analysis reveals roles for candidate phyla and other microbial community members in biogeochemical transformations in oil reservoirs. MBio, 7 (1):e01669–15, 2016..

40. Erik D. Demaine, Felix Reidl, Peter Rossmanith, Fernando Sánchez Villaamil, Somnath Sikdar, and Blair D. Sullivan. Structural sparsity of complex networks: Random graph models and linear algorithms. CoRR, abs/1406.2587, 2014. URL http://arxiv.org/abs/1406.2587.

41. Wojciech Nadara, Marcin Pilipczuk, Roman Rabinovich, Felix Reidl, and Sebastian Siebertz. Empirical evaluation of approximation algorithms for generalized graph coloring and uniform quasi-wideness. In Gianlorenzo D’Angelo, editor, 17th International Symposium on Experimental Algorithms, SEA 2018, June 27-29, 2018, L’Aquila, Italy, volume 103 of LIPIcs, pages 14:1–14:16. Schloss Dagstuhl - Leibniz-Zentrum fuer Informatik, 2018.. URL https://doi.org/10.4230/LIPIcs.SEA.2018.14.

42. Martial Marbouty, Axel Cournac, Jean-François Flot, Hervé Marie-Nelly, Julien Mozziconacci, and Romain Koszul. Metagenomic chromosome conformation capture (meta3c) unveils the div ersity of chromosome organization in microorganisms. eLife, 3, dec 2014.. URL https://doi.org/10.7554/elife.03318.

43. Christopher W. Beitel, Lutz Froenicke, Jenna M. Lang, Ian F. Korf, Richard W. Michelmore, Jonathan A. Eisen, and Aaron E. Darling. Strain- and plasmid-level deconvolution of a synthetic metagenome by sequencing proximity ligation products. PeerJ, 2:e415, may 2014.. URL https://doi.org/10.7717/peerj.415.

44. Migun Shakya et al. Comparative metagenomic and rrna microbial diversity characterization using archaeal and bacterial synthetic communities. Environ. microbiol., 15(6):1882–1899, 2013..

45. Qingpeng Zhang, Sherine Awad, and C. Titus Brown. Crossing the streams: a framework for streaming analysis of short DNA sequencing reads. https://doi.org/10.7287/peerj.preprints.890v1, mar 2015. URL https://doi.org/10.7287/peerj.preprints.890v1.

46. Daniel Standage, Ali yari, Lisa J. Cohen, Michael R. Crusoe, Tim Head, Luiz Irber, Shannon EK Joslin, N. B. Kingsley, Kevin D. Murray, Russell Neches, Camille Scott, Ryan Shean, Sascha Steinbiss, Cait Sydney, and C. Titus Brown. khmer release v2.1: software for biological sequence analysis. The Journal of Open Source Software, 2(15):272, jul 2017.. URL https://doi.org/10.21105/joss.00272.

47. Rayan Chikhi, Antoine Limasset, and Paul Medvedev. Compacting de bruijn graphs from sequencing data quickly and in low memory. Bioinformatics, 32(12):i201–i208, jun 2016.. URL https://doi.org/10.1093/bioinformatics/btw279.

48. J. Koster and S. Rahmann. Snakemake-a scalable bioinformatics workflow engine. Bioinformatics, 28(19):2520–2522, aug 2012.. URL https://doi.org/10.1093/bioinformatics/bts480.

49. Thomas Kluyver, Benjamin Ragan-Kelley, Fernando Pérez, Brian E Granger, Matthias Bus-sonnier, Jonathan Frederic, Kyle Kelley, Jessica B Hamrick, Jason Grout, Sylvain Corlay, et al. Jupyter notebooks-a publishing format for reproducible computational workflows. In ELPUB, pages 87–90, 2016.

50. Stéfan van der Walt, S Chris Colbert, and Gaël Varoquaux. The NumPy array: A structure for efficient numerical computation. Computing in Science & Engineering, 13(2):22–30, mar 2011.. URL https://doi.org/10.1109/mcse.2011.37.

51. John D. Hunter. Matplotlib: A 2d graphics environment. Computing in Science & Engineering, 9(3):90–95, 2007.. URL https://doi.org/10.1109/mcse.2007.55.

52. Wes McKinney. pandas: a foundational python library for data analysis and statistics. Python for High Performance and Scientific Computing, pages 1–9, 2011.

53. Eric Jones, Travis Oliphant, Pearu Peterson, et al. SciPy: Open source scientific tools for Python, 2001 -. URL http://www.scipy.org/. [Online; accessed <today>].

54. Arvind Satyanarayan, Dominik Moritz, Kanit Wongsuphasawat, and Jeffrey Heer. Vega-lite: A grammar of interactive graphics. IEEE Transactions on Visualization and Computer Graphics, 23(1):341–350, jan 2017.. URL https://doi.org/10.1109/tvcg.2016.2599030.

55. Craig A. Stewart, George Turner, Matthew Vaughn, Niall I. Gaffney, Timothy M. Cockerill, Ian Foster, David Hancock, Nirav Merchant, Edwin Skidmore, Daniel Stanzione, James Taylor, and Steven Tuecke. Jetstream. In Proceedings of the 2015 XSEDE Conference on Scientific Advancements Enabled by Enhanced Cyberinfrastructure - XSEDE ‘15. ACM Press, 2015.. URL https://doi.org/10.1145/2792745.2792774.

56. John Towns, Timothy Cockerill, Maytal Dahan, Ian Foster, Kelly Gaither, Andrew Grimshaw, Victor Hazlewood, Scott Lathrop, Dave Lifka, Gregory D. Peterson, Ralph Roskies, J. Ray Scott, and Nancy Wilkens-Diehr. XSEDE: Accelerating scientific discovery Computing in Science & Engineering, 16(5):62–74, sep 2014.. URL https://doi.org/10.1109/mcse.2014.80.

57. Rayan Chikhi, Antoine Limasset, and Paul Medvedev. Compacting de bruijn graphs from sequencing data quickly and in low memory Bioinformatics, 32(12):i201–i208, 2016.

58. C. T Brown, L. Irber, and L. Cohen. dib-lab/sourmash: v1.0. https://doi.org/10.5281/zenodo.153989, September 2016.

59. Sean R Eddy and HMMER Development Team. Hmmer v3.2.1, jun 2018. URL http://hmmer.org/. http://hmmer.org.

60. Robert D. Finn, Penelope Coggill, Ruth Y. Eberhardt, Sean R. Eddy, Jaina Mistry, Alex L. Mitchell, Simon C. Potter, Marco Punta, Matloob Qureshi, Amaia Sangrador-Vegas, Gustavo A. Salazar, John Tate, and Alex Bateman. The pfam protein families database: towardsa more sustainable future. Nucleic Acids Research, 44(D1):D279–D285, dec 2015.. URL https://doi.org/10.1093/nar/gkv1344.

61. K. Katoh and D. M. Standley. MAFFT multiple sequence alignment software version 7: Improvements in performance and usability. Molecular Biology and Evolution, 30(4):772–780, jan 2013.. URL https://doi.org/10.1093/molbev/mst010.

62. Anthony Westbrook, Jordan Ramsdell, Taruna Schuelke, Louisa Normington, R Daniel Bergeron, W Kelley Thomas, and Matthew D MacManes. PALADIN: protein alignment for functional profiling whole metagenome shotgun data. Bioinformatics, 33(10):1473–1478, jan 2017.. URL https://doi.org/10.1093/bioinformatics/btx021.

63. Ryan R. Wick, Mark B. Schultz, Justin Zobel, and Kathryn E. Holt. Bandage: interactive visualization ofde novogenome assemblies: Fig. 1. Bioinformatics, 31(20):3350–3352, jun 2015.. URL https://doi.org/10.1093/bioinformatics/btv383.

64. Minoru Kanehisa, Yoko Sato, and Kanae Morishima. BlastKOALA and GhostKOALA: KEGG tools for functional characterization of genome and metagenome sequences. Journal of Molecular Biology, 428(4):726–731, feb 2016.. URL https://doi.org/10.1016/j.jmb.2015.11.006.

